# PhysiofUS : a tissue-motion based method for heart and breathing rate assessment in neurofunctional ultrasound imaging

**DOI:** 10.1101/2024.09.22.614324

**Authors:** Nicolas Zucker, Samuel Le Meur-Diebolt, Felipe Cybis Pereira, Jerome Baranger, Isabella Hurvitz, Charlie Demené, Bruno Osmanski, Nathalie Ialy-Radio, Valérie Biran, Olivier Baud, Sophie Pezet, Thomas Deffieux, Mickael Tanter

## Abstract

Recent studies have shown growing evidence that brain function is closely synchronised with global physiological parameters. Heart rate is linked to various cognitive processes and previous research has also demonstrated a strong correlation between neuronal activity and breathing. These findings highlight the significance of monitoring these key physiological parameters during neuroimaging as they provide valuable insights into the overall brain function. Today, in neuroimaging, assessing these parameters required additional cumbersome devices or implanted electrodes.

In this work, we performed ultrafast ultrasound imaging both in rodents and human neonates, and we extracted heart and breathing rates from local tissue motion assessed by raw ultrasound data processing. Such ‘PhysiofUS’ automatically select two specific and optimal brain regions with pulsatile tissue signals to monitor such parameters.

We validated the correspondence of these periodic signals with heart and breathing rates assessed using gold-standard electrodes in various conditions in rodents. We also validated Physio-fUS imaging in a clinical environment using conventional ECG.

We show the potential of fUS imaging as an integrative tool for simultaneously monitoring physiological parameters during neurofunctional imaging. Beyond the technological improvement, this innovation could enhance our understanding of the link between breathing, heart rate and neurovascular activity both anesthetised in preclinincal research and clinical functional ultrasound imaging.

## Introduction

Neuronal activity is highly synchronised with empirical physiological parameters such as breathing rate and heart rate. Heart rate variability is linked to autonomic nervous system function and can provide insights into emotional states and stress responses, while respiratory patterns can reflect changes in brain activity related to arousal and alertness. Indeed, heart rate correlates with a wide range of global biological processes, including stress, pain, and sleep.^1-3^ On the other hand, breathing is a fundamental rhythm of brain function, with growing evidence of strong synchronisation between brain activity and breathing rate.^4-6^ Breathing is also implicated in sensorial rhythms and attention modulation.^7-10^ Therefore, these rhythms have been monitored and recorded for years during *in vivo* experiments to obtain quantitative information on the global health status of the subject. Abnormalities in cardiac and respiratory rates are associated with neurological and psychiatric diseases. Irregular heart rate patterns have been observed in patients with epilepsy, anxiety disorders, and depression.^11,12^ Respiratory patterns can also be altered in conditions such as sleep apnea and panic disorder.^13^ Synchronising these physiological parameters with neuroimaging could offer deeper insights into the functional and dysfunctional connections between neuronal processes and cardiac and respiratory rhythms.

Several innovative approaches for monitoring physiological parameters have emerged, particularly for small animal models.^14,15^ These include smart wearable textiles, stretchable electronics, and battery-free implants.^16-18^ However, integrating wireless multiparametric sensors to monitor physiological parameters during neurofunctional imaging remains challenging. It is mainly done at the cost of increased setup complexity, higher costs, and the challenge of synchronising multiple modalities. This is especially relevant to ultrasound neuroimaging of the brain, which, alongside preclinical fMRI, is one of the most widely used imaging modalities in preclinical research.^19^ Indeed, implanted electrodes remain the reference device for assessing breathing and heart rates in most preclinical setups for functional imaging of rodent brains. They can be used to monitor the depth of anaesthesia,^20^ trigger imaging sequences,^21^ and study the influence of drugs.^22,23^ However, these sensors yield strong limitations; they are mainly compatible with anaesthetised rodents and they increase the number of probes and cables used for the experiment. They also increase the clutter on the animal’s head and in some cases, the electrode needles can disturb the rodent’s behaviour. Similarly, in clinical experiments, a significant challenge lies in finding dedicated devices that do not interfere with neuroimaging. Therefore, developing an all-in-one approach for monitoring physiological parameters in both preclinical and clinical imaging alongside functional neuronal activity is crucial. This would allow cost and bulk reduction, automatic synchronisation as well as enabling these parameters to be measured in new experimental situations.

Thanks to the significant increase in frame rate capabilities and sensitivity to blood flow,^24^ ultrasound imaging is able to measure neuronal activity at the whole brain scale.^25,26^ Functional Ultrasound (fUS) imaging can be performed non-invasively in mice and through chronic transparent cranial windows in rats or larger animal models.^27,28^ Combined with intravenous microbubble injections, it can reveal the vascular network and the neurovascular response down to the microscopic range.^29,30^ In the last decade, the acquisition frame rate of ultrasound imaging increased from 50 Hz in conventional imaging to 1000 Hz in ultrafast ultrasound.^31,32^ This increase enabled the tracking of tiny tissue motions induced by fast transient mechanical vibrations, such as shear waves, and has opened up the field of shear wave elastography for cancer and cardiovascular diagnosis.^33^ While Shear Wave Elastography originally relied on the generation of shear waves via the radiation force of ultrasonic focused beams,^34,35^ it is also feasible to image the propagation of natural mechanical waves, such as arterial pulse waves^36,37^ or muscular contractions.^38-40^ Natural shear waves induced by cardiac pulsatility generate physiological mechanical noise, typically ranging between 1 and 50 µm in local displacement, which can be exploited to map the mechanical properties of organs.^41^ In ultrasound localisation microscopy, this estimation of pixel-sized drift from ultrasonic speckle tracking is also applied to improve spatial resolution and correct motion artefacts.^42^ Additionally, this motion detection can be used to extract the cardiac pulsatility on short time scales, allowing synchronisation of ultrasound frames with the cardiac cycle and extraction of systolic and diastolic vascular properties.^43,44^ In other preclinical imaging modalities, such as functional magnetic resonance imaging (fMRI), the link between mechanical motion and physiological parameters has been studied in humans.^45,46^ Specifically, variations in heart rate and respiration can affect the BOLD signal and should be carefully considered for functional connectivity mapping using fMRI.^47^

In this work, we demonstrate that local ultrasonic estimation of brain tissue motion allows reliable detection and quantification of both heart and breathing rates during *in vivo* functional ultrasound imaging in rodents and human neonates, without requiring any additional device for physiological monitoring. The approach, termed “PhysiofUS”, is based on the computation of transient brain tissue motion obtained from Tissue Doppler of raw-IQ data acquired at ultrafast frame rates using fUS imaging. Two very specific regions of interest are then automatically selected in the imaging slice with an original algorithm. Finally, the tissue motion averaged over these two regions yields the heart rate and breathing rate respectively. This automatic fUS-based quantification of physiological parameters was initially performed transcranially in mice and in trepanned rats in both sagittal and coronal planes. The mean absolute percentage error of this measure relative to the gold standard was found to be less than 2% in anaesthetised rodents, demonstrating the high accuracy of this method as an integrated tool for assessing physiological parameters. We further show that the method “PhysiofUS” is robust for tracking breathing and heart rates in non-anaesthetised and freely moving animals, even in the presence of potential motion artefacts. Finally, we transposed this methodology into clinics and showed that ultrasound is able to simultaneously record the brain neurovascular activity and to extract the heart rate of sleeping human neonates by performing tissue motion estimates through their fontanelle. The ultrasound-based estimates of heart rate are derived from fUS raw data and found to match the ECG ground truth, indicating the high translational value of this pipeline in clinics.

## Materials and methods

### Animal experiments

All experiments performed in this study complied with the French and European Community Council Directive of September 22 (2010/63/UE). They were also approved by the local Institutional Animal Care and Ethics Committees (#59, ‘Paris Centre et Sud’). The data collected in this study come from animals from various studies in our laboratory. None of these animals were anaesthetised; or sacrificed for the purpose of our study. These data were acquired during the experiments planned in these different protocols: anaesthetised mice or rats, awake and freely moving rats. The number of animals included, the nature of protocols, and the ethic approval numbers are listed in the *Supplementary Table 1*. The number of animals in our study was kept to the minimum necessary.

Experiments were performed on *N=4* male adult mice (C57BL/6 Rj, age 2-3 months, 25-30 g, from Janvier Labs, France) and *N=13* rats (Sprague Dawley, 325g from Janvier labs, France). See *Supplementary Table 1* for individual details on the animals included. Rats and mice were housed in two and four per cage, respectively, in controlled conditions (22 ± 1°C, 55 ± 10% relative humidity, 12/12 h light/dark cycle), with *ad libitum* access to food and water. Before the beginning of the experiments, the animals were acclimated to the housing conditions for a minimum of one week.

#### Sex as a biological variable

Our study only examined male mice and rats because female animals were not present in our laboratory at the time of the acquisitions. It is unknown whether the findings are relevant for female mice.

#### Anaesthesia

Mice were anaesthetised by intraperitoneal injection of a mixture of Kétamine/Xylazine (100mg/kg, 10mg/kg). Rats were anaesthetised by induction of isoflurane 4%, followed by maintenance (once placed in the stereotaxic frame) at 1·5% (50% O2, 50% air as carrying gas).

For all animals, the depth of anaesthesia was monitored throughout the imaging session from the heart (using ECG electrodes placed subcutaneously, AD Instruments & Labchart) and breathing rates (Spirometer, AD instruments, Labchart). The temperature of the animal was also monitored during the procedures using a rectal probe connected to a heating pad set at 37 °C (Physitemp, Clifton, USA). Eyes of mice were protected using a protective gel (Ocry-gel, TVM, UK).

#### Surgical implantation of ultrasound clear skull prosthesis for head-fixation in behaving rats

As previously reported by M. Matei and colleagues^48^, adult male Sprague Dawley rats aged 7-8 weeks were first put through a week of habituation with the experimenter and before undergoing surgical craniotomy and implantation of an ultrasound-clear prosthesis. Before surgery, anaesthesia was induced by breathing 3·5% isoflurane in a closed space, and by injection of Vétergésic (0·05 mg/kg) and Lurocaine (7 mg/kg). It was maintained with 2·5% isoflurane (50% O2, 50% air as carrying gas). A sagittal skin incision was performed across the posterior part of the head to expose the skull. We excised the parietal and frontal flaps by drilling and gently moving the bone away from the dura mater. The opening exposed the brain from Bregma +4·0 mm to Bregma -9·0 mm, with a maximal width of 14 mm. For rats used in sleep experiments, a stereotactically implanted electrode was secured to the flap’s edge. A prosthetic skull made of polymethylpentene (250 μm thickness, Goodfellow, Huntington UK, goodfellow.com) was sealed in place with acrylic resin (GC Unifast TRAD), and the residual space was filled with saline. In sleep experiments, electrodes were implanted in the animals’ heads to enable monitoring of their current sleep phase. A decision tree based on various electrode measures was used to synchronise fUS data with sleep-scoring phases. A detailed description of the sleep scoring methodology can be found in the work of M. Matei and colleagues^48^.

#### Free exploration session

After at least 10 days of recovery, the animals were fit to undergo a week of habituation, in which the animal was gradually habituated to the room, experimenter, the arena in which it was imaged, and finally the probe-mediated connection to the fUS imager. To perform the 2D imaging, 1 mL of centrifuged ultrasonic gel was applied on top of the chronic window. The ultrasonic probe was positioned and held in place using a 3D-printed magnetic probe holder (in-house design). The animal was then placed inside a 1 x 1 m arena under an infrared camera (to monitor the behaviour) and the data acquisition was started. At the end of the recording session, the window was cleaned from excess ultrasonic gel and the animal was replaced in its home-cage. All rats used in the paper were taken from other experiments being performed in the lab without interfering with the results.

**Supplementary Table 1:**
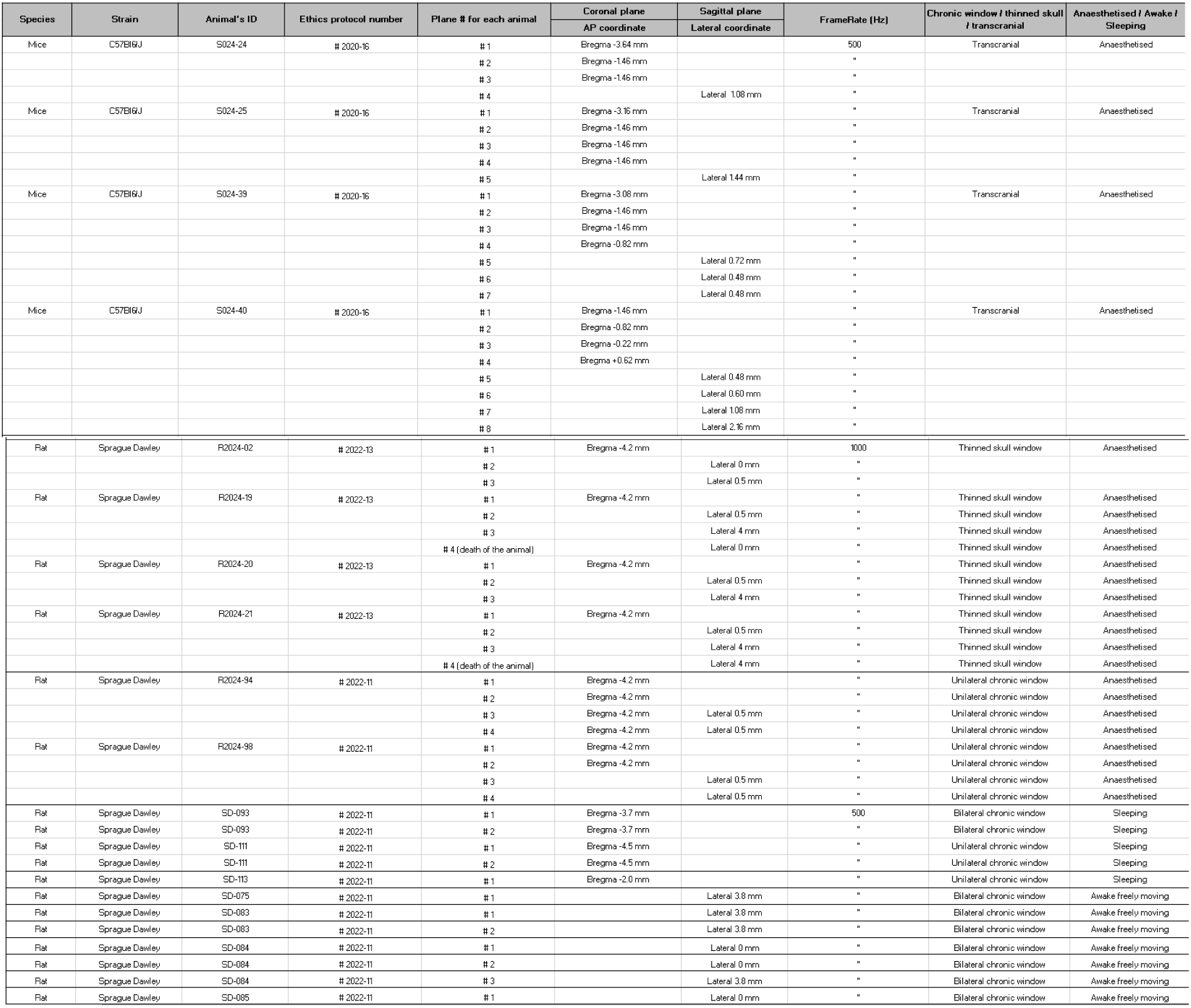
Description of the number of included animals, imaging planes and their associated ethics protocol and reference publication.

#### Preparation of the imaging sessions in rodents

After the animal was anaesthetised, the skin over the skull was shaved with depilatory cream, rinsed several times, and the animal was placed in a stereotaxic frame to ensure the stability of the imaging plane. An echographic gel layer of 1 mm was placed on top of the skull. The ultrasound plane was chosen using linear motors (3 translation + 1 rotation motor) driven by the Iconeus neurofunctional ultrasound imaging scanner (Iconeus, Paris, France). In rats, a linear scan (12 mm span along the anterior-posterior axis, 0·2 mm step) was performed at the beginning of each session using the ‘IcoScan’ software (Iconeus, Paris, France). This scan was used to align the imaging slice with a standard Doppler reference template that was already pre-aligned with the Waxholm rat brain atlas.^49,50^ This allowed automatic and reproducible placement of the probe above the desired coronal and sagittal imaging plane.

#### Ultrasound imaging and sequences

Functional ultrasound imaging acquisitions were performed with a 1D linear transducer (15 MHz central frequency, 128 elements, 110 μm spatial pitch) connected to a functional ultrasound scanner (Iconeus One, Iconeus, Paris, France and Inserm ART Biomedical Ultrasound, Paris France). An ensemble of 11 plane waves tilted from -10° to +10° were fired at a 5·5 kHz pulse repetition frequency, and the backscattered echoes of each group were coherently compounded into 2D images at either 500 Hz or 1000 Hz, depending on the experimental conditions *(Supplementary Table 1)*.

Power Doppler images were computed by applying a clutter filter based on Singular Value decomposition (SVD) of each block of 200 consecutive frames, as reported by Demene et al.^51^ The SVD threshold was fixed to 30. Each pixel of the final power Doppler image was reconstructed on a 110 x 100 μm grid and a slice thickness of 400 μm. Functional ultrasound acquisitions lasted between 2 to 10 min for anaesthetised experiments and 60 min for sleep sessions.

#### Assessment of ground-truth physiological parameters from implanted electrodes

Breathing rate (BR) was monitored by a pillow placed under the animal’s body and connected to the Power Lab system. Heart rate (HR) was measured using 3 electrode needles placed under the skin and connected to the same device. These two signals, acquired at 1000 Hz were recorded and post-processed in MATLAB rather than directly with built-in labchart tools to overcome user dependency thresholds and to increase the final resolution of breathing and heart rate. A 3 Hz Butterworth low-pass filter was applied to the respiratory motion signal and the MATLAB peak detection function^52^ was used to compute the average period of the signals over time. The parameters of this function were adjusted according to the experimental design (rats or mice) due to variations in the shape and frequency of the signals. These parameters are detailed in *Supplementary Table 2*. This pipeline led to final frequencies of 1 Hz for the heart rate and 0·5 Hz for the breathing rate.

**Supplementary Table 2.** Peak detection parameters for the extraction of tissue motion from Powerlab device and fUS imaging Data. Intensity threshold corresponds to the minimum peak heights the function detects. Time between peaks the minimal distance in seconds between detected peaks. The window size and step are corresponding to the time-period selected to compute the average heart and breathing rates. Std peak threshold corresponds to the threshold fixed to select time-periods in which the standard deviation of time between peaks is sufficiently high. Low and high frequency thresholds are the frequency boundaries the algorithm is tracking. In neonates data, a periodogram was used to extract heart rate.

| Application of the peak-detection | Intensity threshold (%) | Time between peaks (sec) | Window size / step (sec) | Std peak threshold | Low frequency thresholds (bpm) | High frequency thresholds (bpm) |
| --- | --- | --- | --- | --- | --- | --- |
| PowerLab Heart Mice | 98 | 0.1 | 2 / 1 | 10 | / | / |
| PowerLab Breathing Mice | 70 | 0.1 | 4 / 2 | 30 | / | / |
| PowerLab Heart Rats | 98 | 0.1 | 2 / 1 | 10 | / | / |
| PowerLab Breathing Mice | 70 | 0.1 | 4 / 2 | 30 | / | / |
| ECG Heart Rate Neonates | Height>0.15 | / | 4 / 2 | / | 120 | 220 |
|  | Intensity / prevalence |  |  |  |  |  |
| fUS Heart Mice | 0.3 0.5 | 0.1 | 2 / 1 | / | 100 | 300 |
| fUS Breathing Mice | 0.3 0.5 | 0.1 | 4 / 2 | / | 200 | 1500 |
| fUS Heart Rats | / | 0.1 | 2 / 1 | / | 40 | 100 |
| fUS Breathing Mice | / | 0.1 | 4 / 2 | / | 200 | 1000 |
| fUS Heart Rate Neonates | / | / | 4 / 2 | / | 120 | 220 |

### Translation to human neonates

Neonatal datasets from a previously published work were reused. All technical details are provided in Baranger *et al*.^53^ Six healthy preterm neonates aged 28+/-2 weeks (post-menstrual age) were included in the study, which was approved by the local ethical board (Robert Debré Hospital, Comité de Protection des Personnes #120601, BELUGA protocol, promoted by INSERM) with strict observance of the World Medical Association Declaration of Helsinki for medical research involving human subjects. All parents gave written and informed consent. fUS datasets were acquired through their fontanelle using a dedicated 6 MHz ultrasound headset connected to a clinical-grade ultrafast ultrasound scanner (Aixplorer, SuperSonic Imagine, France). Due to hardware constraints on the data-transfer rate, fUS acquisitions were discontinuous and constituted of blocks of 352 ultrasound frames acquired at 600 Hz (i.e a 570ms duration) followed by a 430ms pause. One to three 20 min-long fUS acquisitions were obtained during sleep for each neonate. The electroencephalogram (EEG), ECG and breathing were simultaneously recorded and synchronised with fUS acquisition using a Nihon-Kohden EEG-1200 system. The sleep phases such as active or quiet sleep were derived from the EEG tracing by identifying typical patterns (e.g: ‘Trace discontinue’ for quiet sleep^54^). Similar to animal experiments, the heart rate of neonates was derived from ECG tracings with filtering parameters detailed in *Supplementary Table 2*.

### PhysiofUS pipeline and validation

#### Dynamic mapping of local brain tissue motion

Frame-to-frame tissue displacement was evaluated before extracting blood flow signals *(Figure 1.b.)* using the lag-1 correlation of successive 3D (x, z, t) complex in-phase and quadrature signals.^55,56^ The resulting signals were spatially averaged using a 2D spatial filter of width two voxels. A third-order Butterworth temporal filter was then applied with a centre frequency of 20 Hz before smoothing again using a 2D spatial average filter of width two voxels. Tissue motion was finally computed as the phase angle of the autocorrelation signal and averaged over different spatial regions to construct what will be called mean tissue motion (mm) or mean tissue pulsatility (mm/s).

**Figure 1.**
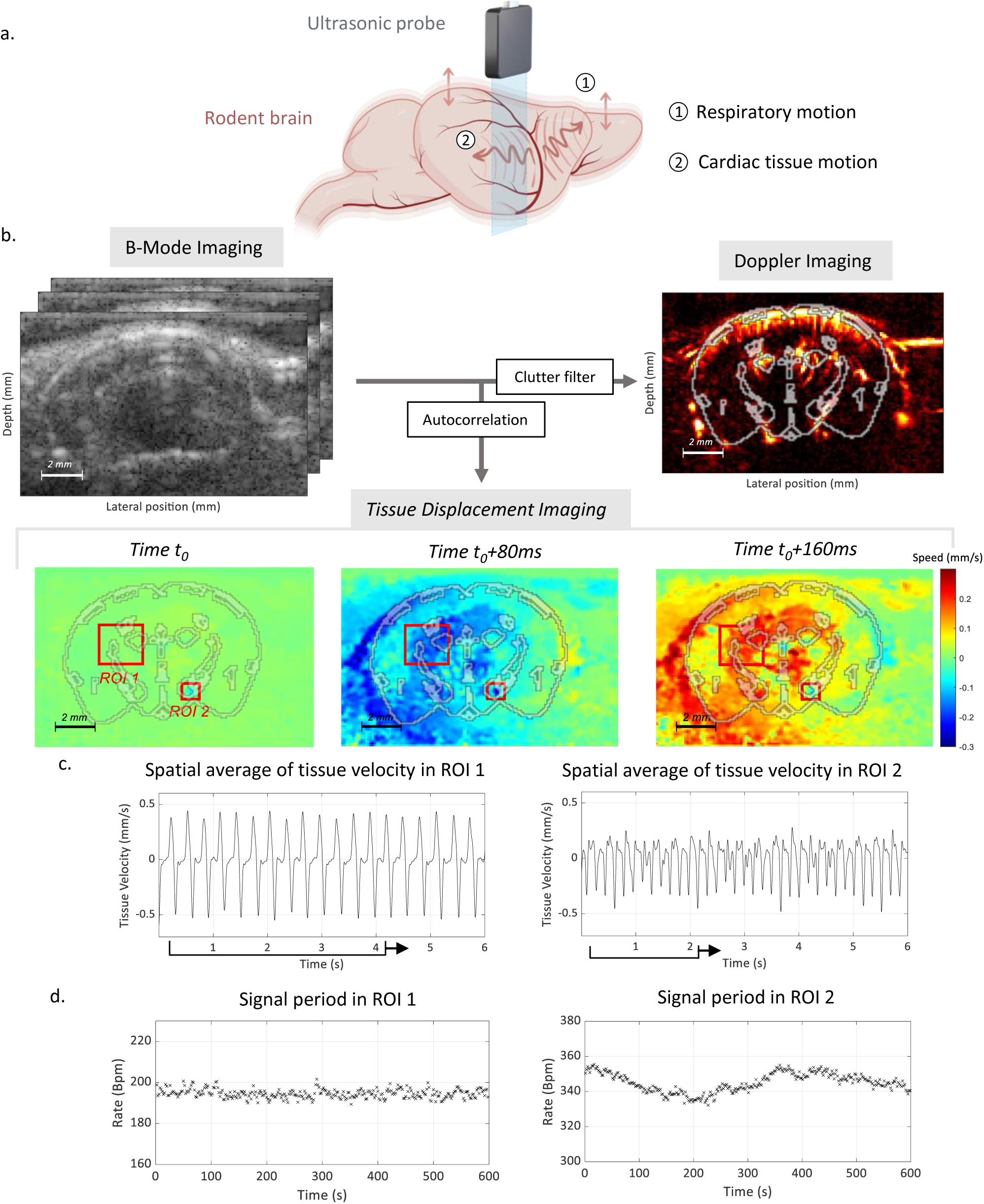
Respiration and heartbeats create periodic tissue motions in the rodent brain detected in fUS. a. Diagram of brain tissue motions being detected by a linear ultrasonic probe. b. B-Mode Imaging, Power-doppler and tissue displacement in functional ultrasound imaging (15Mhz 128 elements) in a coronal acquisition in mice overlaid with an atlas at Bregma - 3·1mm. c. Average tissue velocity in a 6-sec time window in two manually defined region of interests ROI1 and ROI2 shown in b. d. Average periodicity of the tissue velocity using a sliding window of 4-sec for ROI 1 and 2-sec for ROI 2 over 10-mins acquisition.

#### Detection of respiratory movements and heartbeats from local tissue vibrations

In a transcranial 2D fUS imaging session of anaesthetised mice at coronal slice Bregma +2·1 mm, we analysed the mean tissue velocity in two manually defined square regions of interest (*ROI1* 2·0 x 2·0 mm and *ROI2* 0·8 x 0·8 mm). The tissue velocity amplitude was on the order of 1 mm/s. The analysis produced two periodic signals with distinct frequencies *(Figure 1.c)*: a large-scale signal related to breathing *(Figure 1.a*) and another signal related to the rodent’s cardiac pulsatility. In ROI1, the average signal period was 195 bpm (± 2 bpm), while in ROI2, the average period was 345 bpm (± 5 bpm) (*Figure 1.d*). These two frequencies matched the expected breathing and heart rates of anaesthetised mice,^57-59^ and exhibited very low standard deviations. As shown in *Supplementary Figure 2*, random selections of regions of interest led to spatially averaged signals with more complex shapes and varying amplitude and periodicity. These signals were not all reliably related to the expected breathing and heart rates. Thus, the detection of the physiological rhythms depends strongly on the selection of well-defined, plane-specific, and possibly animal-specific regions of interest. Applying singular value decomposition to 3D motion data suggests that this decomposition can efficiently decipher the respective motion caused by cardiac pulsatility and respiratory motion, by leveraging their distinct spatial location and temporal characteristics (*Figure 2, Supplementary Figure 3*).

**Figure 2.**
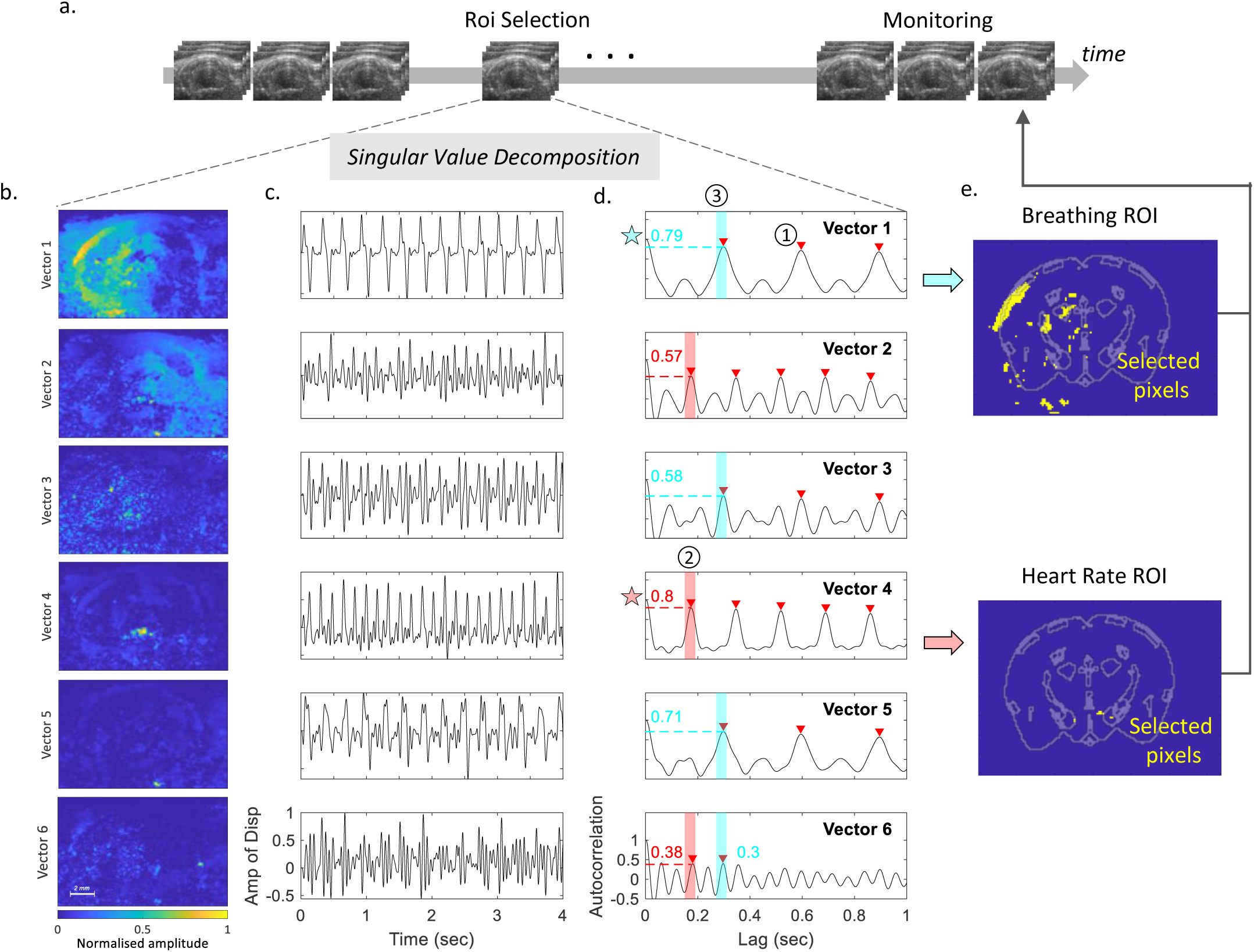
Singular Value Decomposition of tissue motion to identify regions of the brain corresponding to heart and breathing rate. a. 3D frame-to-frame tissue displacement is computed in a 4-sec window to identify the two regions. The signal is processed using space-time singular value decomposition. b. The first six more energetic spatial singular vectors are shown in the left-hand panel. c. The corresponding time vectors as well as their normalised autocorrelation are displayed in the middle panel. One singular vector is selected for the computation of heart and another for breathing rates, the process relies on the : 1. Detection of peaks in the autocorrelation signals. 2. Selection of the most autocorrelated signal for the first peak (corresponding to the HR). 3. Selection of the most autocorrelated signal for the second peak (breathing). Selected eigenvectors are highlighted by a red star for HR and blue star for breathing. e. 2D masks are extracted from the corresponding HR and BR spatial singular vectors, respectively. A mouse brain atlas at Bregma - 3·1 is overlaid.

#### Singular Value Decomposition of tissue motion and automatic selection of singular vectors

The 3D (x, z, t) tissue motion data was calculated as described above, and a Singular Value Decomposition of the corresponding 2D (space, time) Casorati matrix was computed. Time and space vectors corresponding to the six most energetic singular values were retained for further analysis *(Figure 2.b,c)*. While some temporal vectors (such as the second and sixth) did not exhibit any well-defined periodicity, others (such as the first and fourth) matched the signals found in *Figure 1*. The first spatial singular vector, corresponding to a 3·2 Hz periodic signal encompassed a large region of the left brain, while the fourth spatial vector, associated with a 6 Hz periodic signal, was linked to a small region in the centre of the brain. Similar decomposition results, presented in *Supplementary Figure 3*, were obtained from other coronal and sagittal acquisitions in mice and exhibited the same features. A selection process was designed to extract the singular vectors corresponding to heart and breathing rates. The process was based on two assumptions: i). that two among the six most energetic temporal singular vectors pulse at the frequency of the animal’s heart and respiration rates, respectively; ii). the vector corresponding to the respiration rate has a lower frequency than the vector corresponding to the heart rate. These assumptions were verified in almost all animals and experimental designs (*Figure 2, Supplementary Figure 3*). The normalised autocorrelation of each temporal vector was calculated, and the MATLAB peak detection function was used with a minimum peak threshold of 0·3 and a minimum distance between peaks of 0·12 seconds. The temporal positions of these peaks, reflecting the periodicity of the singular vector, were sorted. The first peak position across all vectors corresponded to the highest frequency detected in tissue pulsatility, presumably the heart rate. To mitigate small differences across singular vectors, we defined a temporal window of 0·01 s around the first peak and selected the temporal singular vector that displayed a peak with the highest relative height inside this window. Thus, the selected vector was expected to be closest to the animal’s heart rate. The corresponding spatial singular vector was used to extract the region of interest in which the cardiac pulsatility signal is calculated. The temporal vector assumed to correspond to the breathing rate was selected similarly among the remaining vectors, using the second peak position across all vectors.

In rats, the singular vector identified as breathing was chosen by first applying a 2·5 Hz low-pass temporal filter on temporal singular vectors and then computing the same peak detection algorithm (with a minimum peak distance of 0·5 s). The singular vector corresponding to heart rate was selected among the first six singular vectors time-filtered with a 5 Hz high-pass filter. Peak detection was then applied on the normalised autocorrelation (peak threshold of 0·6, minimum peak height of 0·25, a minimal peak distance of 0·02 s).

#### Automatic region of interest selection for cardiac and respiratory motion extraction

From the previous step, one of the singular vectors marked by a blue star was identified as corresponding to the breathing signal, while another marked by a red star corresponded to the heart rate signal, as shown in *Figure 2.b* and *Supplementary Figure 3*. Next, spatial masks were constructed from the corresponding spatial singular vectors. To extract the breathing rate, we retained 300 pixels with the highest intensities for mice or 1000 pixels with the highest intensities for rats. This threshold was set at 100 pixels in the case of heart rate in rats, whereas in mice, we retained the pixels whose intensity was greater than 80% of the maximum value. These two spatial masks were then used to estimate average tissue pulsatilities, corresponding to respiratory and heart rates. The automatic process of ROI selection for estimation of heart and breathing rates is summarised in *Supplementary Figure 4*.

#### Temporal stability of automatic ROIs selection

To address the stability of the spatial region selection, we split a 10-minute acquisition of an anaesthetised mouse into 150 four-second temporal windows. We then computed the distribution of each singular value across all 150 windows, along with the indices of the singular vectors corresponding to the heart and breathing rates. Finally, we computed 2D masks for the heart and breathing rates in each temporal window and averaged them to compute mean regions of interest (*Supplementary Figure 5)*.

The energy distributions showed low variations among the first 20 singular vectors, with an average standard deviation of 0·4 dB for each singular value. The heart rate signal was consistently identified as the fourth singular vector in 100% of the trials, and the breathing rate signal was identified as the first singular vector in 90% of the trials (*Supplementary Figure 4*). The selected ROIs were largely similar across trials.

#### Assessment of breathing and heart rate based on localised tissue pulsatility periodicity

Tissue motion was computed over time in sub-ensembles of 4-second time windows and spatially averaged in the two regions of interest. A low-pass filter was applied to compute respiratory motion, and a high-pass filter was applied to compute cardiac pulsatility. For mice, we also applied SVD whitening of the signal^60^. The mean period of these signals was computed to extract the heart and breathing rates along the acquisition duration. For the electrode assessment, peak detection was performed on the autocorrelation of the tissue signal with a tenfold interpolation. Parameters used for this detection, such as the sliding window for periods between peaks, are described in *Supplementary Table 2*.

### Comparison with reference electrode measurements

For time synchronisation, ultrasound acquisitions were triggered by the labchart software (Adinstruments). The comparison of the two measures was performed in *N=4* mice covering 15 different coronal planes and 9 sagittal planes, and in *N=6* rats covering 8 coronal planes and 15 sagittal planes (*Supplementary Figures 6 and 7*). A one-second delay was applied to the electrode signal to account for the time required by the ultrasound scanner to start the acquisition. Differences in heart rate and breathing rate measured using the fUS and ground truth methods were assessed at the same time points. These curves were computed across four experimental designs (coronal and sagittal, rats and mice) and fitted using the linear model *fUS = a*Electrodes + b*. Regression coefficients and coefficients of determination were computed. The difference time curves between the two approaches were calculated, and the time averages of the absolute values of these differences for each experimental session were grouped into four categories: coronal mice, sagittal mice, coronal rats, and sagittal rats. To determine the average expected variation of the fUS measure relative to the ground truth, we computed a 95% confidence interval using a Wilcoxon two-tailed signed rank test. We also quantified heart rate variations in two additional sessions in which the animal was progressively dying (*Supplementary Figure 8*).

#### Evaluation of the computation time of the pipeline

To assess the usability of the pipeline for real-time monitoring with a functional ultrasound scanner, we evaluated the performance of a MATLAB implementation on a 6-core computer (AMD Ryzen 5 5600X). We measured the time needed for the initialisation part in *N=24* acquisitions in mice and *N=25* acquisitions in rats. We then extracted a 2-second signal of cardiac pulsatility and respiratory motion from a random temporal window. These computations did not take into account the time required to load data into MATLAB, as this would be negligible during real-time monitoring (Supplementary Figure 9).

#### Processing of rat sleep experiments

We applied the fUS pipeline to pre-existing data used in the paper of Marta Matei.^48^ The variations in whole-brain Cerebral Blood Volume (CBV), obtained as the spatial average of the power Doppler signal, were computed and displayed alongside varying heart and breathing rates to emphasise the capability of extracting physiological parameters as additional measures during functional brain imaging. The commonly defined sleep stages (Active Wake, Quiet Wake, Non-REM Sleep, and REM Sleep) were overlaid on the plots as colour patches. The pipeline’s success rate was computed as the proportion of data points where physiological measurements met quality thresholds and fell within the heart rate range of 250 – 400 bpm and the breathing rate range of 40 – 100 bpm.

#### Quantification of the variation of physiological parameters during sleep phases

We first evaluated variations in HR at the onset of sleep. In two acquisitions in which the animal was awake at the start of the session before falling asleep, we extracted fUS-based measures of HR in the first 50 points (100 s) and compared it to the distribution of HR measured 5 min later. A two sample t-test was performed between the two distributions. We then quantified variations in breathing rate between sleep phases in *N=5* acquisitions in 3 animal sessions lasting between 50 and 60 minutes. We identified transitions from sleep states and calculated the average value of the physiological parameter during the 30 seconds before and after each transition. A two-sample t-test was then performed to compare the means of breathing and heart rates between sleep states across acquisitions.

#### Processing of rodents awake freely moving acquisitions

The fUS pipeline for assessing physiological parameters was applied to imaging sessions in freely moving rats. The size of the selected region of interest from the spatial singular vectors was reduced to the tenth greatest pixel of the selected vector. We synchronised the video recording with fUS data and displayed the variations of the animals’ speed, whole brain cerebral blood volume, and heart rate computed throughout the acquisition. We finally computed the success rate for each acquisition session, defined as the percentage of time during which a measure of the heart rate was successfully extracted between 200 bpm and 450 bpm.

#### Application of the pipeline to sleeping neonates

Brain tissue motions were derived in neonates acquisition with the same method used in awake and freely moving rodents (*Figure 5.d*). The heart rate of the patient was assumed to range from 120 bpm to 250 bpm, which are typical values for this population. As ultrasound acquisitions blocks were not continuous, a Lomb-Scargle periodogram^61^ was used to evaluate the heart rate along fUS acquisitions, with computational parameters described in Supplementary Table 2. HR measurements from fUS and ECG were synchronised in *N=6* neonates and compared using a linear model: *fUS = a*Electrodes + b*.

## Results

### Local tissue motions in some brain regions are linked to breathing and heart rates

While performing brain imaging in rodents, large-scale movements due to the animal’s breathing and more localised higher frequency movements due to heartbeat can be detected from the speckle-noise correlation of raw ultrasound data *(Figure 1.a,b,c*). The cardiac pulse wave propagating through arteries induces small shear mechanical motions in tissues surrounding the arterial wall.^62,63^ Applying singular value decomposition to 3D (x, z, t) motion data can effectively differentiate the respective motions caused by heart and breathing rates, by leveraging their distinct spatial and temporal characteristics (*Figure 2, Supplementary Figure 3,4*). We therefore, proposed an automatic method to select two regions of interest where tissue motion signals reflect respiratory motion and cardiac pulsatility. The temporal window used for this operation has minimal impact on the selected regions (*Supplementary Figure 5*). Finally, the averaged estimates of the signal periods give access to heart and breathing rates (*Figure 1.d*).

### Validation of heart-rate and breathing-rate measurements with electrodes

To evaluate the robustness and accuracy of our approach, we synchronised functional ultrasound measurements with measurements from implanted electrodes. The experimental setup for these recordings, where the fUS acquisition is triggered by the PowerLab system is presented in *Figure 3.a*. Examples of the synchronised extraction of physiological parameters in transcranial mice and rats are presented in *Figure 3.b. and 3.c*. In the mice acquisition, the average temporal values respectively for heart and breathing rates were 363·3 and 208·7 bpm for fUS assessment, and 364·1 and 208·7 bpm for electrodes measurements. Additional comparisons are presented in *Supplementary Figure 6 and 7*. The variation of heart and breathing rates between the two approaches, quantified by the linear model *fUS=a*Electrodes+b* in three experimental setups, yielded regression coefficients ranging from 0·92 to 1·08 and linear regression coefficients between 0·87 and 0·98 (*Figure 3. d*).

**Figure 3.**
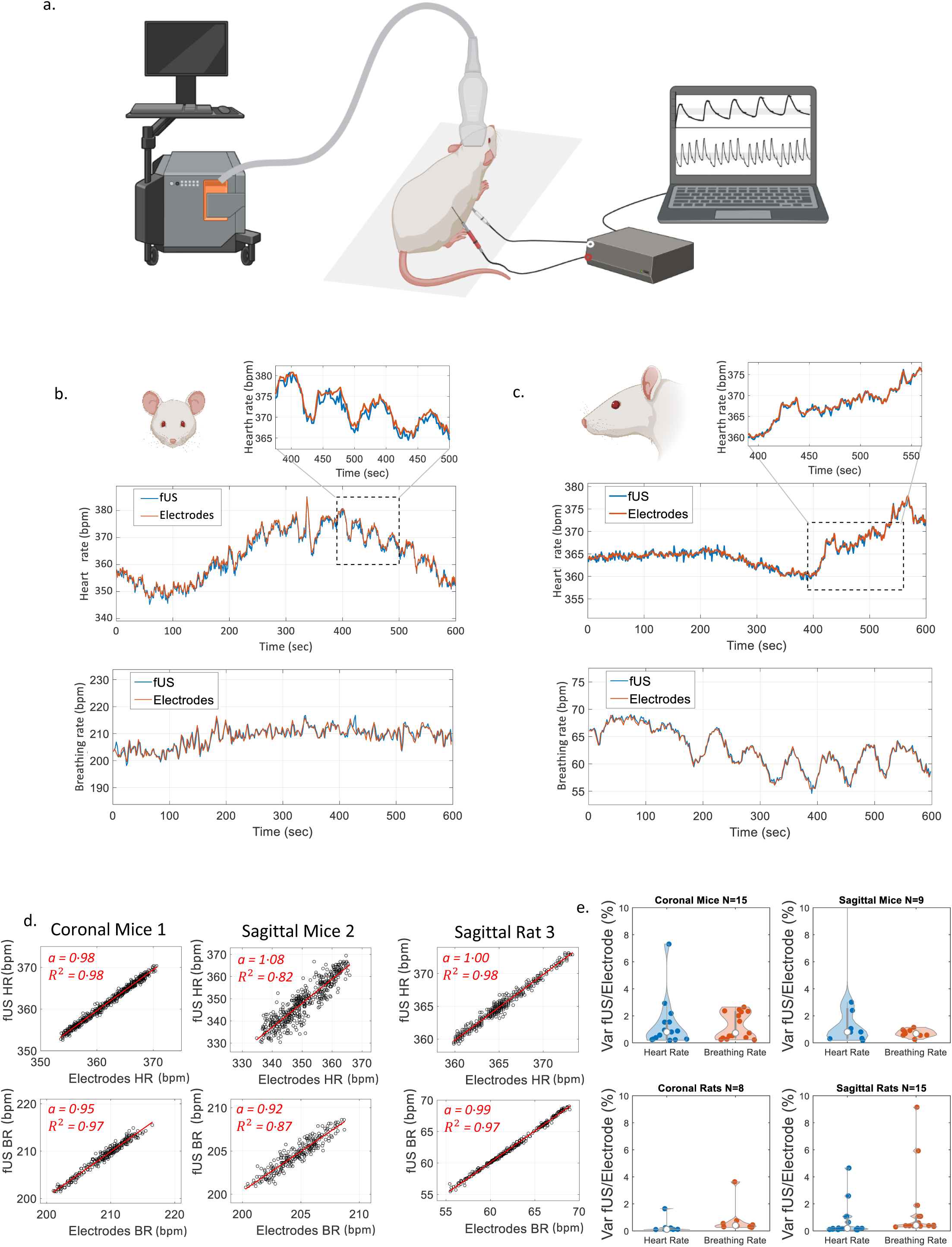
Synchronisation of fUS and reference electrodes assessment of respiration and heart rate in anesthetized mice and rats. a. Schematic diagram of the experimental design used to synchronize electrodes and ultrasound acquisition. The ultrafast ultrasound scanner is triggered by the PowerLab device. Figure created with BioRender.com. b. Superposition of heart rate (top) and breathing rate (bottom) assessed with electrodes and fUS in a coronal mouse during a ten-minute acquisition. A zoom into a two-minute time window is shown. c. Superposition of the heart rate (top) and breathing rate (bottom) assessed with electrodes and fUS in a sagittal plane in a rat during a ten-min acquisition. A zoom into a 2-minute time window is shown. d. Quantification of the variations of heart rate and breathing rate assessed with the two approaches. Curve modelling of the slope *fUS = a*Electrodes + b* for HR and BR in 3 acquisitions. Regression coefficient and coefficient of determination are computed and shown. e. Quantification of the mean percentage of the absolute difference between functional ultrasound and electrodes in the 4 experimental conditions: coronal mice, sagittal mice, coronal rats and sagittal rats (error bars are showing first and last quartiles).

We compared the average absolute variation between fUS and electrode assessments during 10-minutes acquisitions in various experimental conditions, grouped into four categories: coronal mice, sagittal mice, coronal rats, and sagittal rats (*Figure 3.e*). The median error across all mice acquisitions is 0·95% ([0·56%-1·7%] 95% CI) for heart rate and 0·94% ([0·57%-1·44%] 95% CI) for breathing rate in mice. In rats, the median error is 0·22% ([0·14%-0·64%] 95% CI) for heart rate and 0·55% ([0·38%-1·16%] 95% CI) for breathing rate.

We applied the same approach in two experimental sessions in rats where the animals’ progression towards death led to significant heart rate changes of 20% and 65%. Modelling the fUS-obtained heart rate against the electrode assessment yielded a linear regression coefficient of 0·999. (*Supplementary Figure 8*).

#### Towards real-time assessment of physiological parameters with a fUS scanner

While the previous extraction of breathing and heart rate was done on previously acquired data, the processing can be optimised for real-time applications. We have outlined a pipeline, illustrated in *Supplementary Figure 9*, that could be implemented in an ultrafast ultrasound scanner to assess physiological parameters in real-time. To verify the feasibility of this approach, we estimated the computation time for the various steps of the pipeline.

The average computation time for the initialisation was 5·3 s (± 0·76 s) for mice and 25 s (± 1 s) for rats. Following this, the tissue motion field was computed in 2-second sub-blocks, and the signal was averaged in the two defined regions of interest. The time required for this operation was 1·06 s (± 0·07 s) and 1·21 s (± 0·06 s) respectively. Consequently, the overall duration of the measurement is on the order of the temporal resolution of the fUS acquisitions, allowing for live display of these parameters during experiments.

#### Estimation of physiological parameters in behaving rats

After validating the method on anaesthetised animals, which allowed for direct comparison with electrodes, we tested whether we could apply the same pipeline to behaving rats. Specifically, we conducted two experimental scenarios: one with a rat sleeping in a cage while implanted electrodes monitored its sleep stages, and another with a rat moving freely in an arena.

### Estimation of heart and breathing rates during sleep stages

We applied the method to a set of *N=5* acquisitions (3 animals) in which trepanned rats implanted with a chronic window were habituated to sleep with an attached ultrasound probe *(Figure 4.a)*. We extracted the physiological parameters and whole-brain cerebral blood volume during the 1-hour acquisitions *(Figure 4.b, Supplementary Figure 10)*. We successfully measured heart rate in 41% of the time points and breathing rate in 44% of the time points. We quantified variations in physiological parameters across sleep phases and detected a decrease in heart rate at the onset of sleep *(Figure 4.c)* and its increase during awakening from 288 bpm to 321 bpm *Figure 4.d*. Finally, we computed variations of breathing rate during *N=19* transitions from rapid-eye-movement episodes to non-REM-sleep, finding an decrease from 83 bpm (*std = 8 bpm*) during REM to 75 bpm (*std = 8 bpm*) during Non-REM (two-sample *t-* test p<0.01).

**Figure 4.**
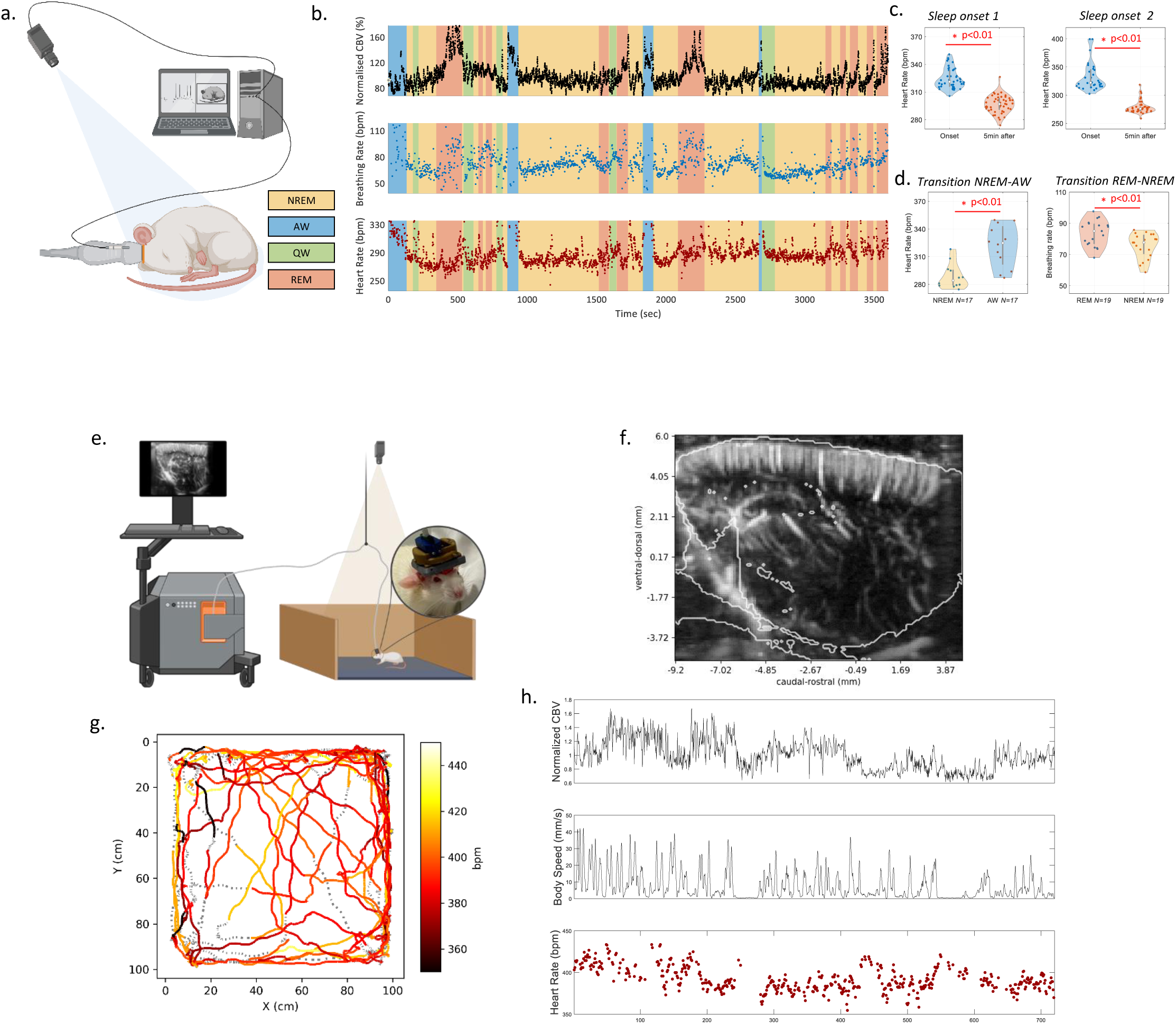
Synchronised functional ultrasound imaging of brain activity and physiological parameters extraction in behaving rats. a. Schematic of the experimental design used for fUS imaging of sleep in rats. b. Synchronised assessment of whole brain cerebral blood volume, heart rate and respiratory rate during a one-hour ultrasound acquisition. c. Quantification of the decrease of heart rate at the onset of sleep in two acquisitions (error bars are showing first and last quartiles). d. Quantification of the variation of heart rate at the transition from NREM to AW and variation of breathing rate at the transition of REM to NREM e. Diagram of experimental design for fUS imaging of neuronal activity in a rodent moving freely in an arena. f. Power-doppler sagittal slice image overlaid with the Waxholm rat atlas at the lateral position -3.8 mm from sagittal median plane. g. Plot of the animal’s displacements in the 1m by 1m arena during the 12-minute acquisition period. The color of the curve corresponds to the measured heart rate, a dashed line symbolises that no heart rate was measure. h. Synchronise variations of the cerebral blood volume, the body speed of the animal detected by a camera and the measured heart rate with functional ultrasound.

### Estimation of heart and breathing rates in a freely moving rat

We applied the method to an experimental design in which a rat moves freely in an arena while undergoing fUS imaging and motion tracking *(Figure 4.d).* The resulting sagittal imaging plane is depicted in *Figure 4.e* overlaid with the Waxholm rat atlas.^50^ We measured the heart rate of the animal throughout the 10-minute acquisition and synchronised it with the animal’s cerebral blood volume and speed *(Figure 4.g)*. We successfully measured the heart rate in 60% of the temporal points and found a mean heart rate of 388 bpm with a standard deviation of 34 bpm. The motion coverage of the animal, with colour indicating heart rate, is displayed in Figure 4.f. The same extraction obtained in another animal shows that the tissue speed signal was stable, showing strong periodicity across different time windows *(Supplementary Figure 11)*. We further evaluated the heart rate estimator in *N=4* animals (*N=7 sessions*) and obtained reproducible results. The success rate for heart rate measurements during each session was found to be 69% ± 29% in this animal group.

### Synchronised functional ultrasound imaging and heart rate assessment in human neonates

In neonates, the PhysiofUS pipeline detected specific regions of the imaging slice in which tissue pulsatility presented periodic oscillations close to the one detected with the ECG (*Figure 5.a,b,c,d,e,f*). Temporal variations of the resulting heart rate extracted from fUS data and synchronous heart rate computed from the ECG are overlaid in Figure 5.g-h, and *Supplementary Figure 12*, in both quiet and active sleep phases. In both subjects 1 and 2, the heart rate was successfully measured with fUS in 100% of time points, with an average absolute variation of 1·0%. Fitting a linear model to all heart rate measures obtained with fUS against all ECG measures in a total of *N=6* neonates yielded a linear regression coefficient of 0·98 and a coefficient of determination of 0·93, for heart-rate spanning a wide range of [130 - 180] bpm (Figure 5.i).

**Figure 5.**
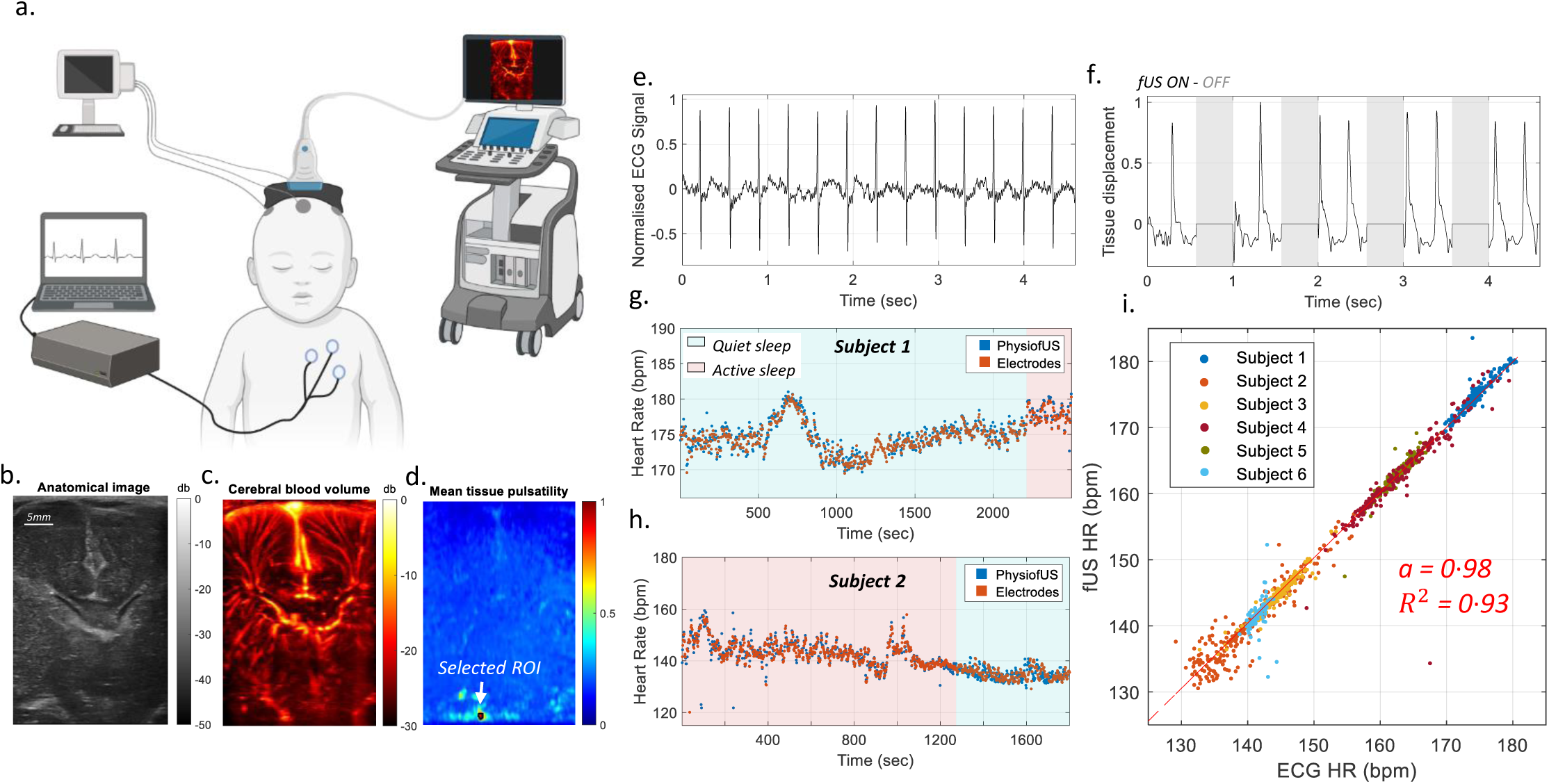
Synchronised functional ultrasound imaging of brain activity and physiological parameters extraction in neonates. a. Schematic of the experimental design used for simultaneous fUS imaging and ECG measurments. b. Anatomical B-Mode image in 2D fUS imaging. c. Power Doppler image computed as the mean intensity of filtrated IQ. d. Square root of the average temporal tissue pulsatility over 1-sec of fUS acquisition. The ROI selected by the pipeline in which the tissue pulsatility is averaged over time is represented. e. Normalised ECG signal used to compute the heart rate of neonates. f. Average tissue pulsatility in the region of interest over a 4.5 temporal windows. Time periods during which no fUS datas are acquired are highlighted with gray patchs. g.h Superposition of heart rate assessed with ECG and fUS in two neonate with a sampling frequency is 0.025Hz, distinct sleep phases are represented by color patches. i. Quantification of the variations of heart assessed with the two approaches in 6 subjects, for clarity 1/3 of the temporal points are displayed. Curve modelling of the slope *fUS = a*Electrodes + b* for HR and BR. Regression coefficient and coefficient of determination are computed is shown.

## Discussion

In this work, we demonstrate that heart and breathing rates can be simultaneously evaluated alongside neurofunctional ultrasound imaging of brain activity in preclinical and clinical acquisitions. In most of the experimental setups, heart and breathing rates can be extracted from brain tissue motion signals. The animal’s breathing creates large-scale motion in the brain, while the heartbeat induces more localised periodic motion in the vicinity of large vessels. We show that it is possible to identify regions of interest specific to heart and breathing rates using space-time singular value decomposition of ultrasound-based tissue motion. This process could be integrated into an automatic pipeline that extracts heart and breathing rates throughout the acquisition. This approach was validated by comparing these values with gold standard reference electrodes. In both anaesthetised rats and mice, and across multiple planes and imaging sessions, the heart and breathing rate measurements from the ultrasound method differed from the reference electrodes by less than 2%. Additionally, we extracted heart and breathing rates in behaving, non-anaesthetised rats during sleep conditions and free exploration in arenas. Finally we showed that heart rate can be extracted simultaneously to fUS in human neonates with a 1% deviation to the ECG gold-standard approach, in conditions where the heart rate of neonates were spanning over a wide range of values. Together, these results suggest that fUS can be used as an all-in-one approach to measure physiological parameters during clinical and preclinical functional ultrasound imaging of the brain.

Singular value decomposition efficiently separates signals into both temporal and spatial components with high energies. Since breathing and heart rates induce tissue motions with high energy along both time and space, SVD is a robust method to extract these physiological measures from tissue motion signals. Tissue motions induced by breathing were found to encompass large areas of the imaged slices and were consistently associated with the first singular vector, i.e. the most energetic vector. In contrast, heartbeat-induced motions were localised in smaller areas from the deepest regions of the brain, corresponding to large cerebral arteries where cardiac pulsatility affects vessel walls. The absence of these vessels in the imaging slices of some ultrasound sessions may hinder the measurement of heart rhythms. Despite this, the proposed ultrasound-based pipeline generally provides accurate estimates of mean heart and breathing rates and reveals important periodic oscillations in these rhythms.^64^

The combination of neurofunctional ultrasound with physiological parameter monitoring holds great potential. In behavioural experiments, our approach uses portable ultrasonic probes mounted on the animal’s head, offering a fully integrated, all-ultrasound technology for both neuroimaging and physiological monitoring. This setup enables observation of animals’ interactions with their environment, other animals, and their internal rhythms. While assessing physiological parameters during sleep phases, we observed an increase in breathing rate during REM sleep, consistent with the literature.^65,66^ Additionally, we detected a decrease in both breathing and heart rates at the onset of sleep. Notably, these two rhythms exhibited greater variability during REM Sleep. We also detected some physiological responses to sleep-state changes in neonates, with an increase in heart rate in the active-wake state.^67^

In active-wake states and in freely moving rodents, locomotion can introduce potential motion artefacts between the probe and the skull or within brain tissues.^28^ These artefacts disrupt tissue motion estimation and can prevent accurate measurements of small tissue motion variations caused by breathing rate and heartbeat during periods of high activity. Interestingly, although heart rate oscillates during locomotion sessions, it tends to increase after the animal reaches peak speeds. This could be due to a lag between physical activity and the heart rate increase or exertive activities such as grooming that the animal engages in without moving.^68^

In a broader perspective, our findings suggest that brain tissue rhythms detected in tissue motion during fUS acquisitions^42,44,69^ accurately reflect the underlying physiological parameters. The implications of this work are significant. First, the simultaneous extraction of heart and breathing rates during any fUS acquisition, without the need for a specific ultrasonic acquisition sequence, paves the way for systematic integration of these physiological parameters into fUS neuroimaging analysis. Given the complex relationship between physiological parameters and neuronal activity,^4,7,9^ this technique could help elucidate the connections between physiological rhythms and brain connectomics, neurovascular coupling, brain states, or circadian rhythms.^70,71^

Secondly, the low computational cost and the compatibility with common ultrasound sequences should allow for live implementation in functional ultrasound imaging scanners. This integration would facilitate live monitoring of physiological parameters during fUS experiments in both clinical and preclinical imaging, reducing the need for additional sensors and simplifying the setup while ensuring precise synchronisation. Extracting physiological parameters during distinct brain states, such as sleep phases, could provide new insights into fundamental processes like NREM and REM sleep,^71-75^ as well as behaviours such as spatial navigation and social interactions.^76-78^ Furthermore, this work paves the way for a better understanding of the effects of drug treatments on both neuronal activity and physiological parameters in preclinical models, with potential translation to humans.^22,23,79^

The method developed in this study shows strong potential for translation to clinical ultrasound imaging. Given the growing applications of fUS in human and newborn imaging,^80,81^ the precision of our approach in sleeping neonates paves the way for more comprehensive assessments of heart rate in neonates, allowing a simplification of the experimental setup and an automatic synchronisation. It is also likely that brain tissue pulsatility carries in itself characteristics of the cardiac rhythm and could be analysed to detect heart rate arrhythmia or other cardiac pathologies.

However, this approach also has significant limitations. First, the regions of interest used to average the tissue motion were selected based on singular value decomposition and estimation of frequency content; we did not evaluate the feasibility of the extraction in more complex cases where heart and breathing rates might overlap. Additionally, our analysis pipeline required parameter tuning specific to the experimental designs studied. A more generalised pipeline could be developed using larger training datasets and incorporating machine learning algorithms. This would result in a more flexible and robust pipeline, particularly for addressing motion artefacts in behaving animals. In some experimental conditions, such as when acquisition blocks were not continuous, the pipeline was not able to extract the breathing rate: it remains unclear whether the underlying brain motion was still present in the data. Finally, further work should involve validating these results in behaving rats during free exploration using gold-standard wireless sensors to assess heart and breathing rates.

Future research avenues should explore applying the proposed method to other commonly used fUS acquisition setups, such as multi-slice acquisitions,^82^ head-fixed mice,^83^ 3D volumetric imaging in mice and rats^84^ or other clinical imaging systems. Additionally, further investigation is needed to establish whether the pipeline remains compatible with the use of injected contrast agents, such as microbubbles, which are increasingly used in brain vasculature imaging.^29,30,81^ Research should also extend to different experimental paradigms such as ischemic or hemorrhagic stroke, functional activation, functional connectivity or drug-induced functional changes.

## Abbreviations

HR: Heart rate
BR: Breathing rate
fUS: Functional ultrasound
CBV: Cerebral Blood Volume
bpm: beats per minute

## Acknowledgments

This study was supported by the European Research Council under the European Union’s Seventh Framework Program (FP/2007-2013) / ERC Grant Agreement n° 311025 and by the Fondation Bettencourt-Schueller under the program “Physics for Medicine”. We acknowledge the ART (Technological Research Accelerator) biomedical ultrasound program of INSERM.

---

**Supplementary Figure 1:**
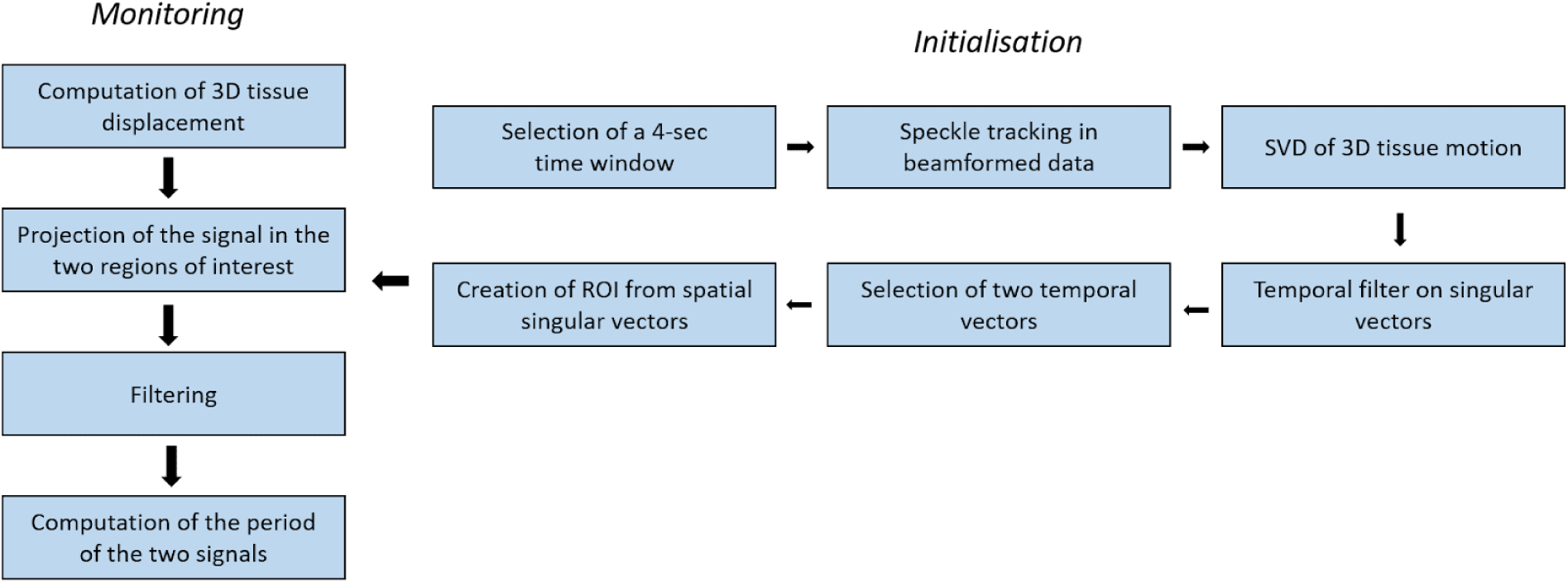
Diagram of the processing steps for ROI selections for physiological parameters estimation.

**Supplementary figure 2.**
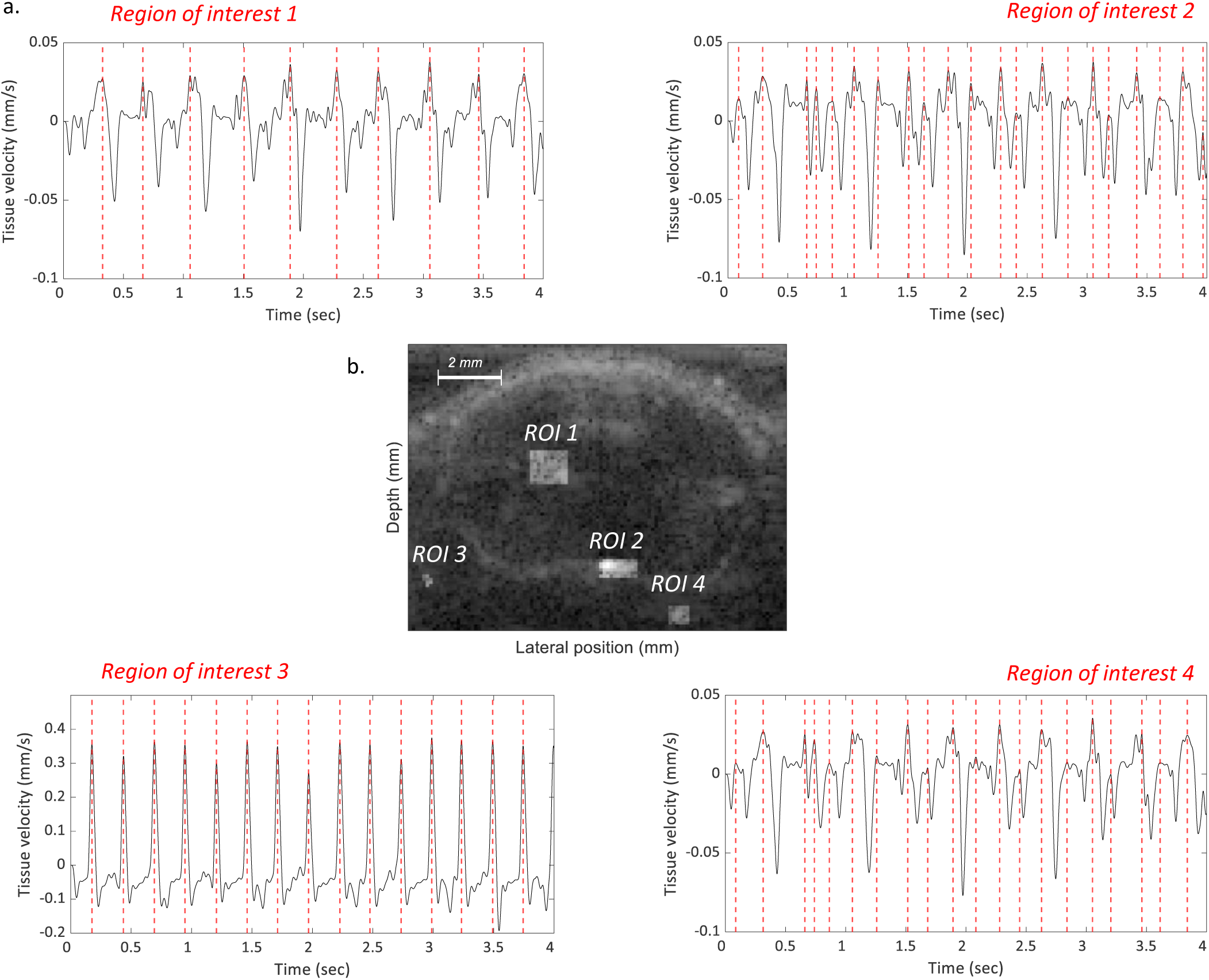
The periodicity of pulsatility in brain tissue depends on the region in which it is averaged. a. **Ultrafast B-mode imaging** averaged over 200 frames at 500Hz overlaid with 4 manually selected regions of interest (ROI). b. **Spatial averaging of tissue velocity in the four regions of interest** over a 4-second time window. Peak detection was applied and displayed by vertical red dashed lines to show periodicity.

**Supplementary figure 3.**
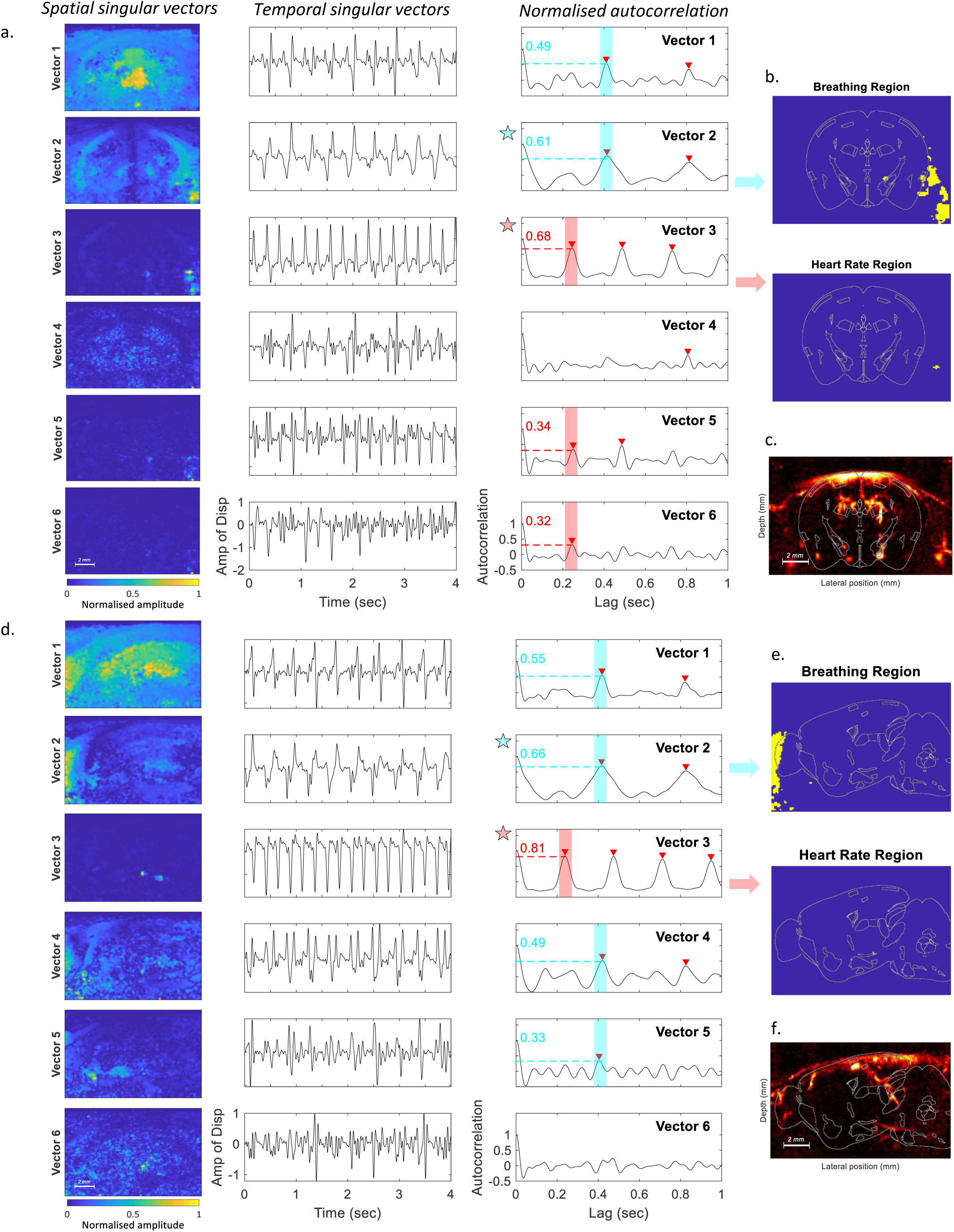
SVD decomposition of tissue velocity and selection of the two regions of interest for breathing and heart rate in a coronal and sagittal mouse. a,d. **Space-time SVD decomposition** and selection of the singular vector corresponding to heart rate and breathing rate. **The selected vector is highlighted** by a red star for heart rate and a blue star for breathing rate. b,e. **Construction of two masks from selected singular vectors**. c,e. 2D **Power Doppler slice corresponding to the acquisition plane** in a coronal (Bregma -1·4mm) and a sagittal (Sagittal-median +2 mm) anesthetized mice mouse.

**Supplementary figure 4.**
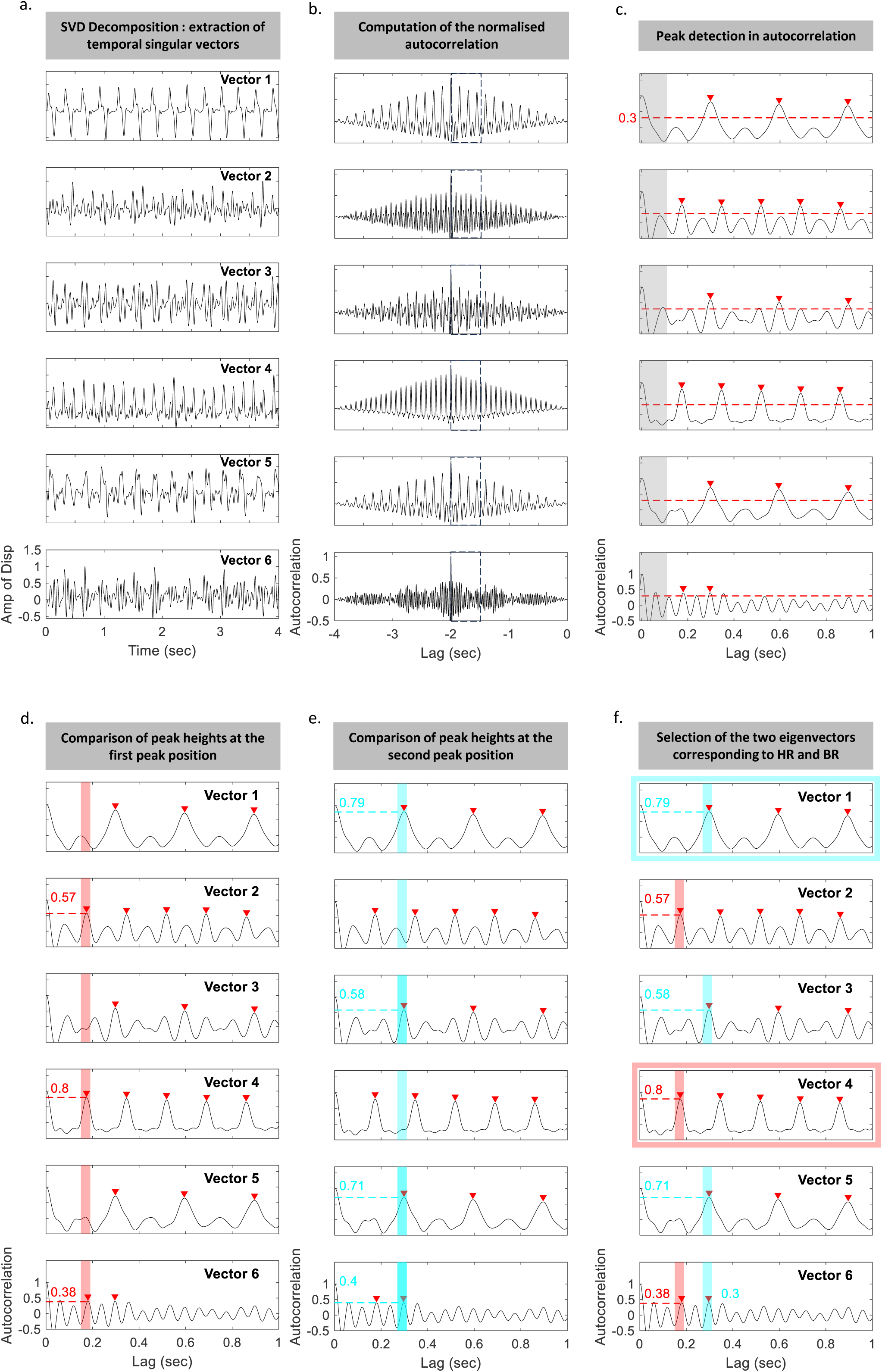
Descriptive diagram of the step-by-step selection of the two singular vectors for extraction of the respiration region of interest and the heart rate region of interest. a. **The six first more energetic temporal vectors are retained**. b**. Normalised autocorrelation of time vectors** is displayed. c. A Matlab **peak detection algorithm** is applied on the extracted region of the autocorrelation (lag greater than 0.1 seconds, 500Hz), represented as a gray patch. The intensity treshold, represented by a red dashed line is fixed to 0.3. Detected peaks are marked with a red arrow. d. **The first time position of the peaks is selected** and the relative heights of the autocorrelation are compared in a temporal window around the first peak (red patch). e. **The second time position of the peaks is selected** and the relative heights of the autocorrelation are compared in a time window around the second peak (blue patch). f. **The two singular vectors selected for heart rate and breathing rate are framed in blue and red.**

**Supplementary figure 5.**
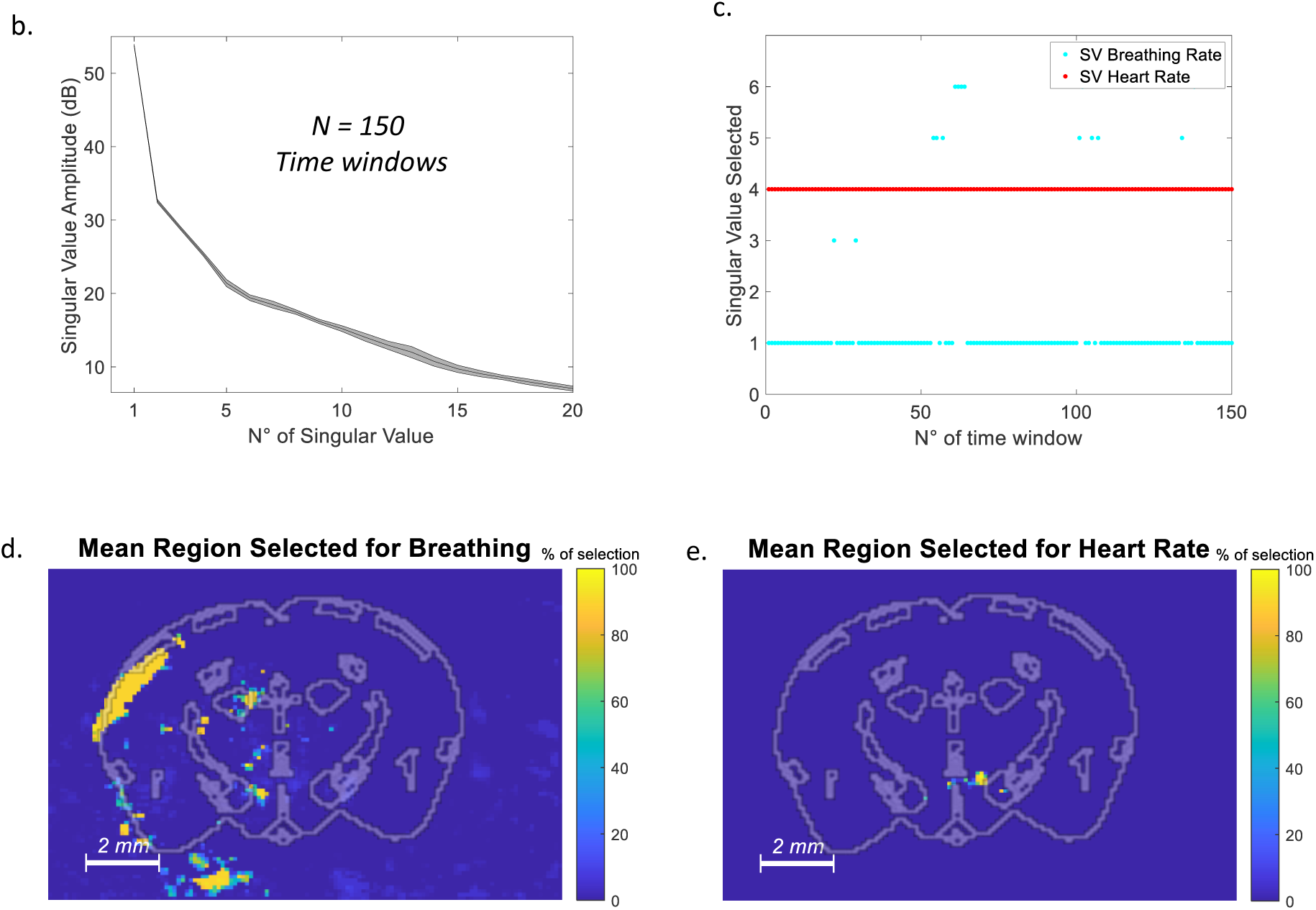
Repetition of the extraction of singular vectors in multiple temporal windows in an anesthetized mice (Coronal Bregma 2.1mm). **a. Singular values amplitude distribution** for the 150 temporal windows of 4sec making up the 10min aquisition. **b. Singular value selected for the heart and breathing rate** in the various temporal windows. c) d) **Time average of the region selected** for respiration (left panel) and heart rate (right panel) by the pipeline in the 150 time windows overlaid with a mouse brain Atlas at Bregma 2.1mm.

**Supplementary figure 6.**
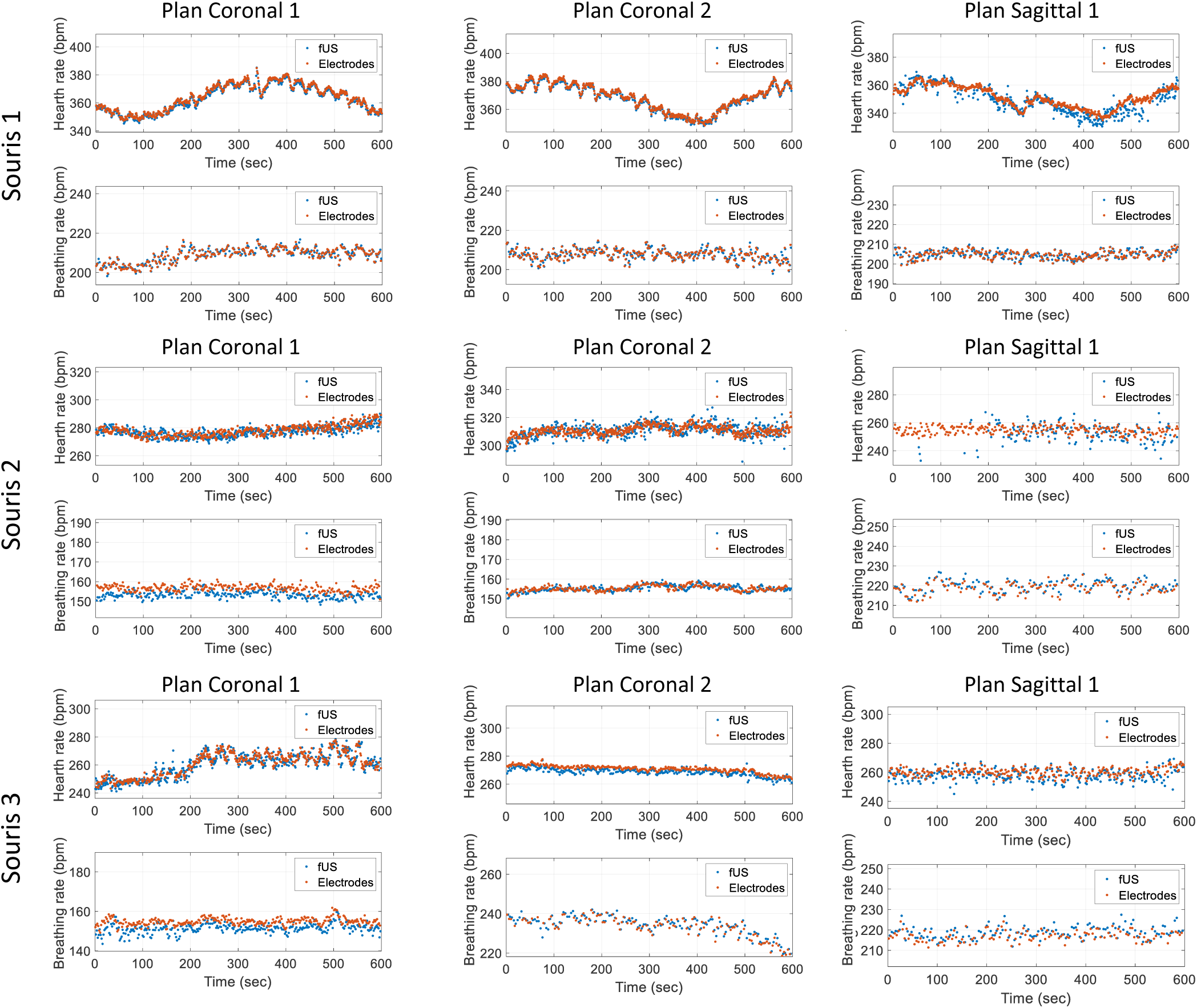
Additional comparison of HR and BR extracted with fUS and with electrodes in 3 mice. For each mice the two mueasures are overlaid in two coronal planes and a sagittal plane.

**Supplementary figure 7.**
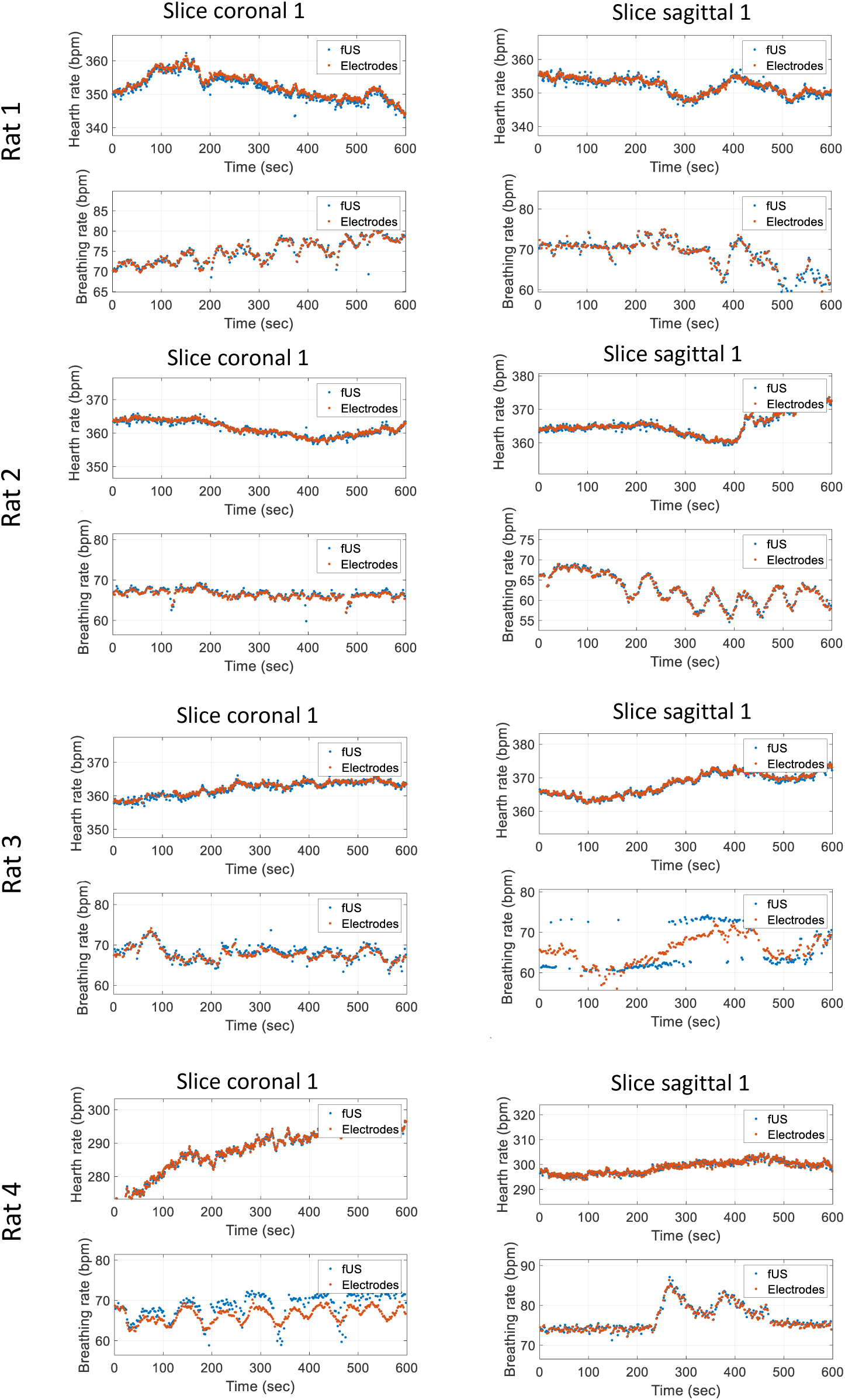
Additional comparison of HR and BR extracted with fUS and with electrodes in 4 rats in one coronal plane and one sagittal plane.

**Supplementary figure 8.**
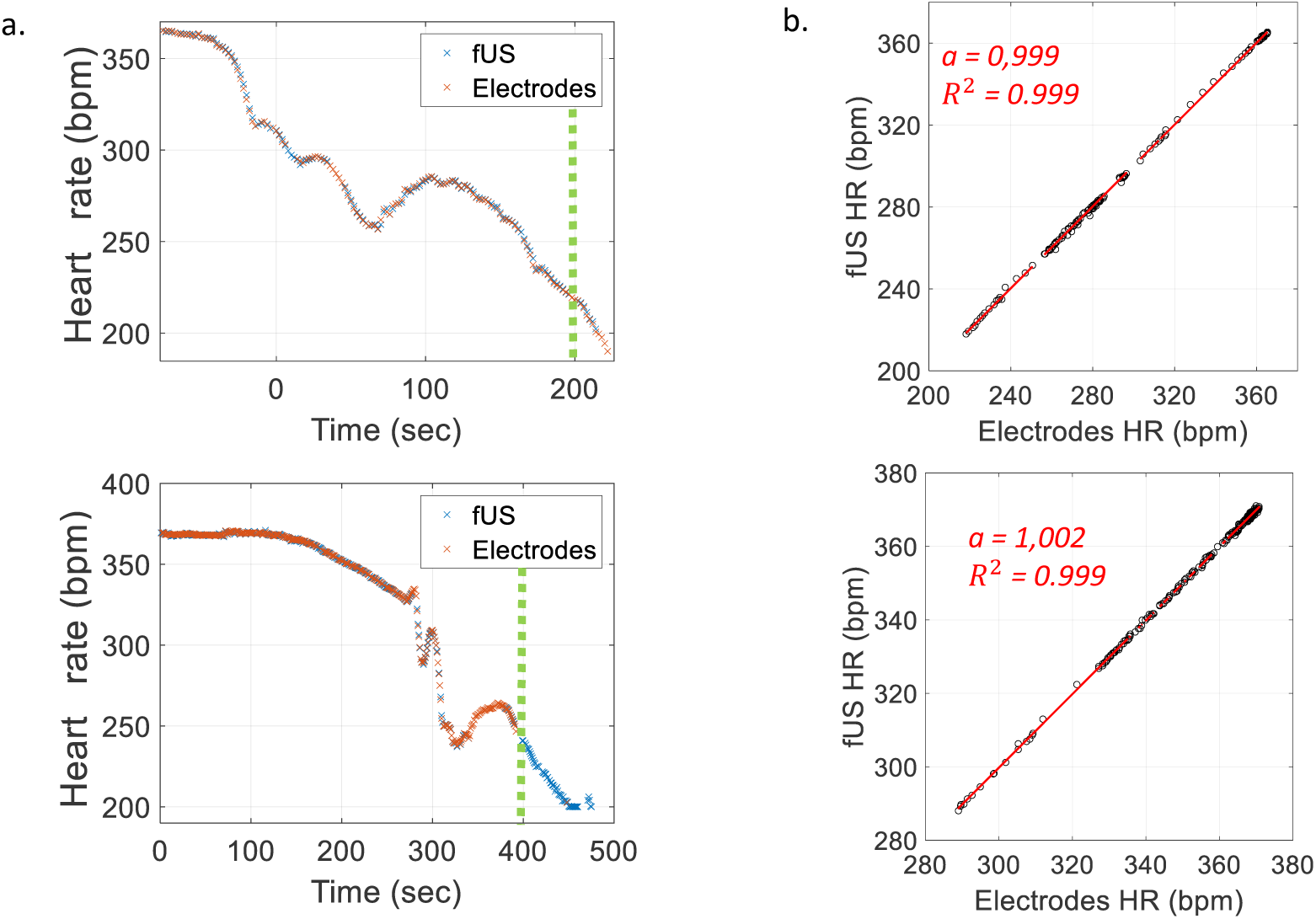
Monitoring of an animal’s heart rate during death using ultrasound and electrodes. a. **Superposition of heart rate assessed with electrodes and fUS** in a sagittal plane two rats that are dying due to the injection of b**. Quantification of the variations** of Heart Rate assessed by the two approaches for the first points of the curve. This limit is represented by the green dashed line. Curve modelling of the slope *fUS = a*Electrodes + b* for HR and BR in 2 acquisitions. Regression coefficient and coefficient of determination are computed and shown. Heart rate values higher than 400 bpm were considered as artefacts (3·6% for the left panel and 1·6% of points for the right one are rejected).

**Supplementary figure 9.**
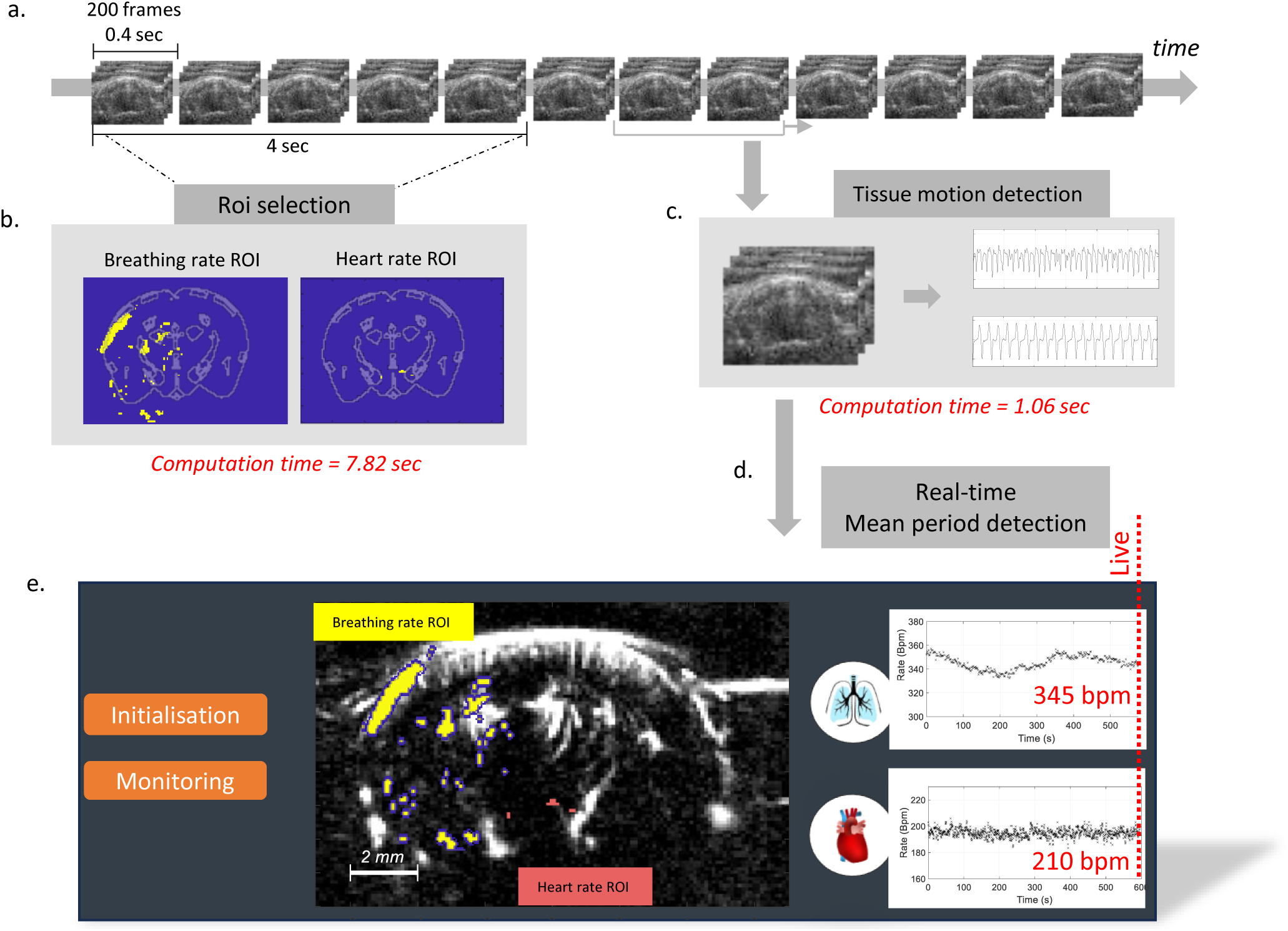
Schematic diagram of the implementation of real-time assessment of heart rate and respiratory rate in an ultrafast ultrasound imaging system. a. **A 10sec time-window is selected** (2000 frames at 500Hz). b. **Breathing rate and heart rate regions of interest are computed** with the pipeline and displayed overlaid with the mouse Atlas at bregma -2.1mm. The average time required for this initialisation is 5.3 seconds in mice. c. **Cardiac pulsatility signal and respiratory motion are computed** as the spatial average of the tissue velocity signal in the two previously defined regions. The time needed for this processing is 1.06 sec d. A **peak-detection algorithm is applied** and the mean period between peaks in displayed. e) **Diagram of the possible screen of an ultrafast scanner** in which the pipeline is implemented, allowing to aquire simultaneously the brain activity (Power-doppler middle-panel) and physiological parameters (right-hand pannel). The process requires an initialisation step and a monitoring one.

**Supplementary figure 10.**
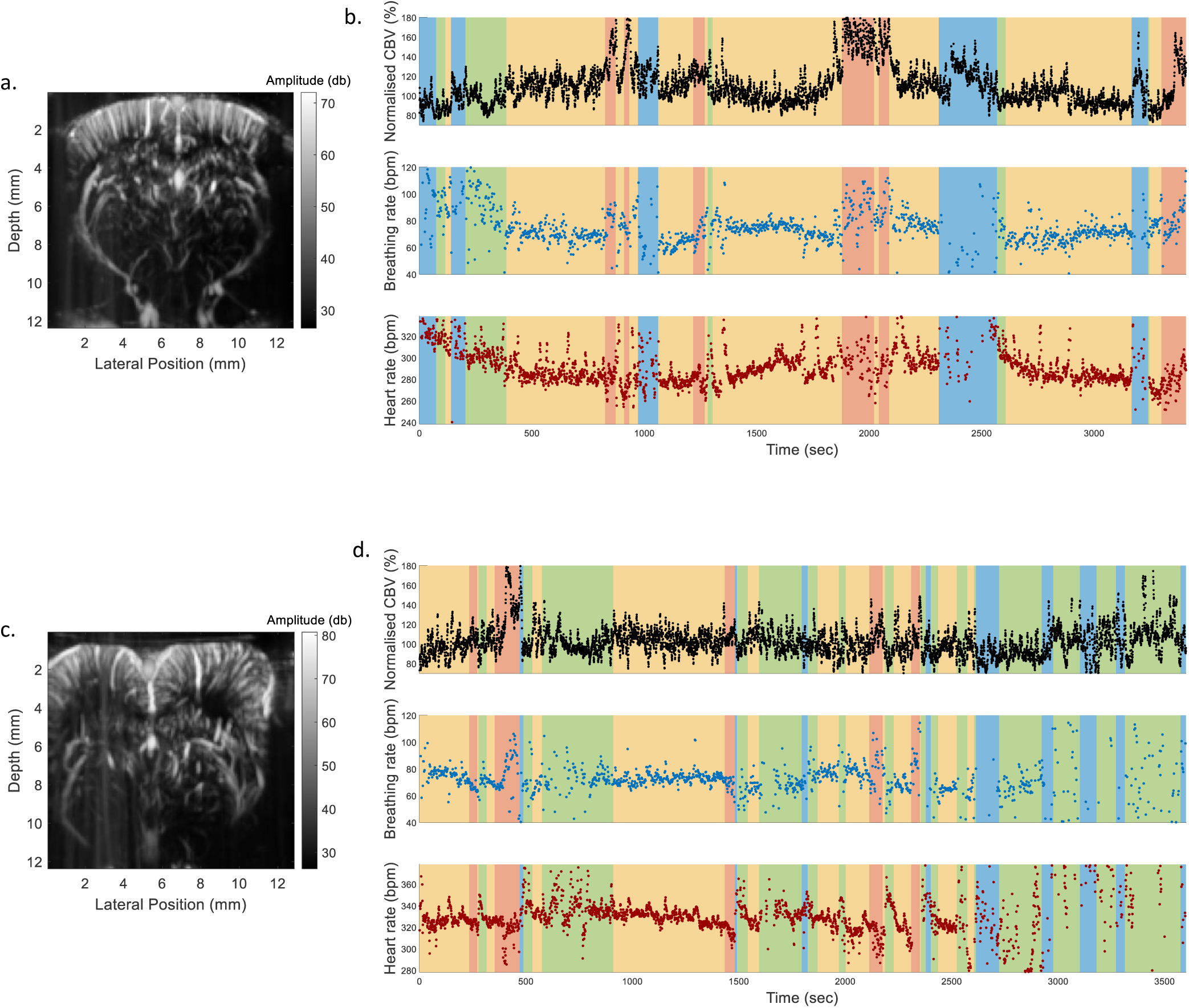
Synchronized assessment of heart rate and respiratory rate during ultrasound imaging of brain activity during a sleep session in two coronal rats acquisition. a,c**. 2D Power-doppler images** of the coronal slice of the rat at Bregma -3,7mm (top) and Bregma 1,8mm (bottom). b,d. **Variation of the whole-brain normalised CBV** assessed by averaging the power-doppler image over time, **with heart and breathing rate assessed with the PhysiofUS pipeline**.

**Supplementary figure 11.**
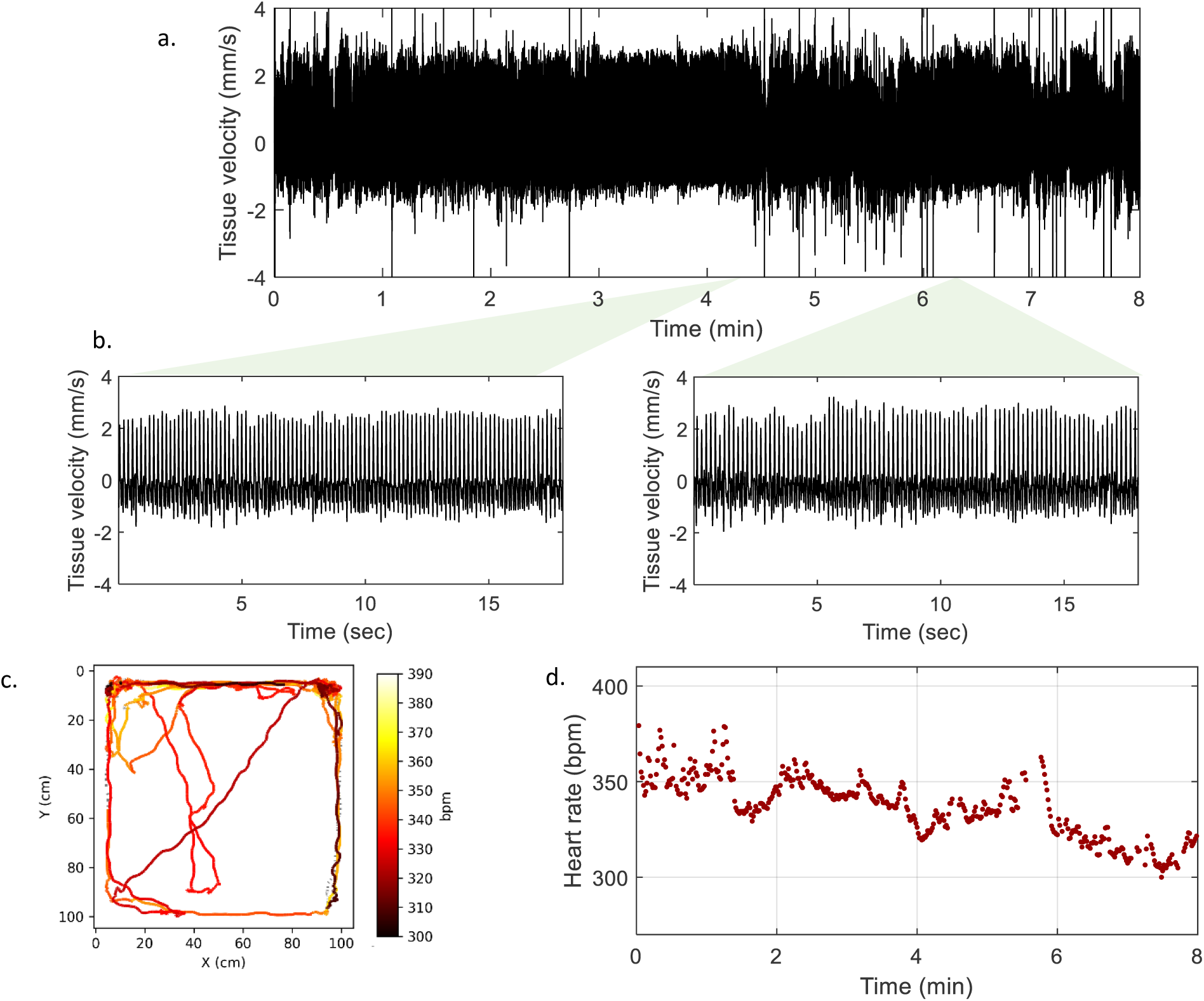
Variations of the speed of tissue in a sagittal rat freely moving in an arena. a. **Average speed of tissue** in the heart rate region of interest during the height minutes of acquisition. b. **Temporal zoom in two temporal regions** of tewenty seconds. c. Plot of the **animal’s displacements in the 1m by 1m arena** during the 8-minute acquisition period. The color of the curve corresponds to the measured heart rate, a dashed line symbolises that no heart rate was measure. c. **Heart rate variation** computed as the average time between peaks in the tissue velocity signal. In this session, the approach allowed to extract heart rate in 88% of the time points.

**Supplementary figure 12.**
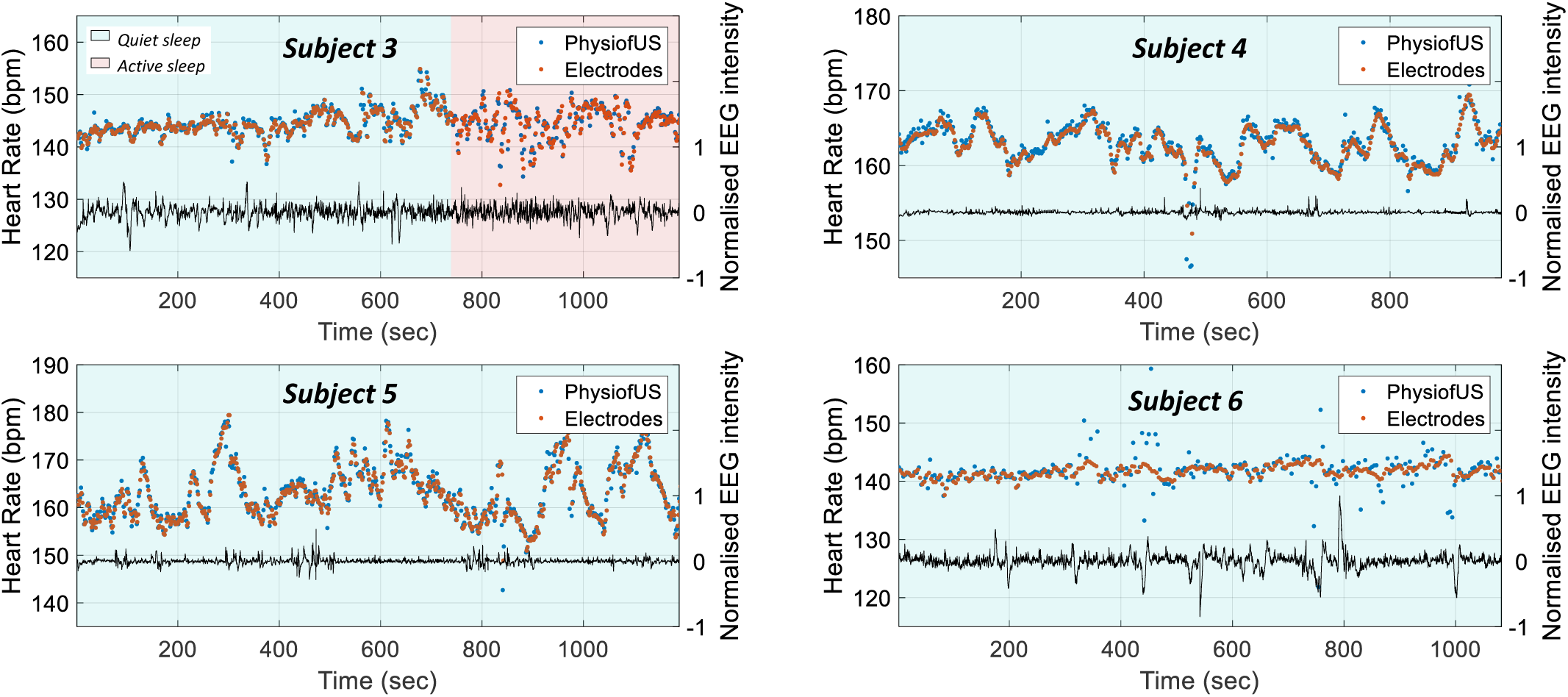
Synchronised functional ultrasound imaging of brain activity and physiological parameters assesment in human neonates. The superposition of heart rate assessed with ECG and fUS in four neonate with a sampling frequency is 0.025Hz is presented, the distinct sleep phases are represented by color patches. Normalised variations of an EEG chanel is plot on the same graph.

